# Extensive High-Order Complexes within SARS-CoV-2 Proteome Revealed by Compartmentalization-Aided Interaction Screening

**DOI:** 10.1101/2020.12.26.424422

**Authors:** Weifan Xu, Gaofeng Pei, Hongrui Liu, Jing Wang, Pilong Li

## Abstract

Bearing the largest single-stranded RNA genome in nature, SARS-CoV-2 utilizes sophisticated replication/transcription complexes (RTCs), mainly composed of a network of nonstructural proteins and nucleocapsid protein, to establish efficient infection. Here, we developed an innovative interaction screening strategy based on phase separation *in cellulo*, namely **<u>co</u>**mpartmentalization of **<u>p</u>**rotein-protein **<u>i</u>**nteractions in **<u>c</u>**ells (CoPIC). Utilizing CoPIC screening, we mapped the interaction network among RTC-related viral proteins. We identified a total of 47 binary interactions among 14 proteins governing replication, discontinuous transcription, and translation of coronaviruses. Further exploration via CoPIC led to the discovery of extensive ternary complexes composed of these components, which infer potential higher-order complexes. Taken together, our results present an efficient, and robust interaction screening strategy, and indicate the existence of a complex interaction network among RTC-related factors, thus opening up new opportunities to understand SARS-CoV-2 biology and develop therapeutic interventions for COVID-19.

## Introduction

The current pandemic of COVID-19 (Coronavirus Disease-2019), a respiratory disease leading to more than 55 million confirmed cases and 1.3 million fatalities globally (WHO, 2020), is caused by SARS-CoV-2, an enveloped, positive-sense, single-stranded RNA betacoronavirus of the family *Coronaviridae* (Zhu et al., 2020). SARS-CoV-2 possesses a remarkably large RNA genome, with a genome size of approximately 30-kb, flanked by 5’ and 3’ untranslated regions. The genomic RNA at the 5’ end features the partly overlapping open reading frames (ORF) 1a and 1ab, that occupy two-thirds of the capped and polyadenylated genome (Lu et al., 2020; Tan et al., 2005). The expression of ORF 1ab requires a ribosomal frameshift near the end of ORF1a. In this manner, SARS-CoV-2 genome translation yields the large replicase polyproteins pp1a and pp1ab (Abduljalil and Abduljalil, 2020; Perlman and Netland, 2009). Pp1a and pp1ab go through extensive autoproteolytic processing, mediated by two ORF1a-encoded proteases and ultimately generating 16 nonstructural proteins (Nsps), which make up the primary constituents of the replication/transcription complexes (RTCs). These include a variety of key enzymes functioning for viral RNA synthesis, i.e. the primase-Nsp8 (Kirchdoerfer and Ward, 2019; Konkolova et al., 2020); the RdRp-Nsp12 (Cheng et al., 2005; Shannon et al., 2020); the helicase-Nsp13 (Jang et al., 2020; Lee et al., 2010); the exoribonuclease-Nsp14 (Minskaia et al., 2006) and the endoribonuclease NendoU-Nsp15 (Bhardwaj et al., 2004); several multi-spanning membrane proteins (Nsp3, Nsp4 and Nsp6) that presumably contribute to the formation of double-membrane vesicles (DMVs) (Cottam et al., 2011; Kanjanahaluethai et al., 2007; Oostra et al., 2007); proteases for polyprotein cleavage (Nsp5) (Stobart et al., 2013), and the accessory modulating factors hijacking host defense (Nsp1 and Nsp2) (Antonin et al., 2020; Cornillez-Ty et al., 2009). For the 3’ one-third of the genome, ORFs that encode four structural proteins and nine accessory proteins are transcribed to form a nested set of subgenomic mRNAs (Abduljalil and Abduljalil, 2020).

The synthesis of SARS-CoV-2 RNA within host cells depends on the efficient and correct assembly of RTCs under the coordinated cascade of transcription and replication. This process takes place in intracellular DMVs (Klein et al., 2020; Snijder et al., 2020), which are actually remodeled to serve as a platform for protein-protein interactions (PPIs) and provide a favorable environment for replication. Although SARS-CoV-2 Nsps (e.g., Nsp7, Nsp8, Nsp12, and Nsp13) are associated with the viral RTCs during infection (Yan et al., 2020b), the molecular details of which Nsps and how they come together to form the viral RTCs are not clear. Notably, it is also reported that nucleocapsid (N) protein functions as the only structural protein shuttling inside/outside RTCs and facilitating the transcription and replication of coronavirus RNA (Cong et al., 2020; Surjit and Lal, 2008). Recently released structural information revealed the interaction of the C-terminal domain of N protein with the pore factor of DMVs (Khan et al., 2020). As a critical step toward understanding this process, it is necessary to investigate the interaction network among Nsps. Information to date in this regard has been scarce, with only a limited number of interactions identified (Pan et al., 2008; von Brunn et al., 2007).

Although multiple methods have been developed to detect PPIs, few are suited for high throughput protein interaction analysis in cells. The most commonly used methods for investigating PPIs of viruses including coronaviruses (He et al., 2004; Pan et al., 2008; von Brunn et al., 2007), yeast two-hybrid (Y2H) screening and coimmunoprecipitation (co-IP), have their own limitations and both suffer severe false-positive and false-negative results (Berggård et al., 2007; Miernyk and Thelen, 2008; Phizicky and Fields, 1995). In addition, co-IP is unfriendly to large-scale screening. To alleviate these drawbacks, here we developed a novel method to study PPIs in living cells, namely **C**ompartmentalization of **<u>P</u>**rotein-protein **<u>I</u>**nteractions in **<u>C</u>**ells (CoPIC). Essentially, CoPIC is the adapted version of a phase separation-based method for PPIs *in vitro* in our previous study, called Phase-separated **<u>C</u>**ondensate-aided **<u>E</u>**nrichment of **<u>B</u>**iomolecular **<u>I</u>**nteractions in **<u>T</u>**est tubes (CEBIT) (Zhou et al., 2020). One major requirement for CEBIT is the availability of large quantities of well-behaved purified materials, which is difficult to achieve for intrinsically ill-behaved subjects such as large dynamic protein complexes. Correspondingly in CoPIC, one interaction partner is fused with the scaffolds that drive the formation of phase-separated condensates, and the other partner (named the client) is fused with a fluorescent protein, and can be recruited to the condensate via specific interactions in cells. Overall, CoPIC emerges as a simple, sensitive, and efficient method to investigate PPI in mammalian cells.

To elucidate molecular mechanisms of viral replication and transcription, there is a need to systematically examine possible intraviral interactions. Although relatively unexplored among the virological studies, intraviral interactions have significant and unexpected effects on host range, transmissibility, immunopathology, and therapeutic effectiveness (Dao et al., 2020; Häuser et al., 2012; Lee et al., 2011; Yin et al., 2019). We, therefore, established an interaction map for RTC-related viral proteins by CoPIC matrix screening and identified a total of 47 binary interactions among 14 proteins governing replication, discontinuous transcription, and translation of coronaviruses. Moreover, comprehensive high-order complexes within the SARS-CoV-2 proteome were also comprehensively deduced. Overall, increased awareness of intraviral interactions as described here is a necessary step for achieving a better understanding of SARS-CoV-2 and thus providing more novel and effective therapeutic opportunities.

## Results

### Establishment of robust scaffolds for phase separation in cells

The design strategy of CoPIC is as follows: i) a system (called scaffold) capable of forming membraneless phase-separated compartments in cells is generated; ii) one component of the PPI of interest (POI) is fused with the scaffold and hence is enriched in the compartments; and iii) the recruitment of another one component (called client) into the compartments serves as the readout for the interaction (**Figure 1A**). For simplicity, we chose phase separation-prone low complexity domains (LCDs) as potential scaffolds to form the compartmental hubs for interactions (Malinovska et al., 2013; Wang et al., 2018; Weber and Brangwynne, 2012). To this end, we generated a series of expression vectors with a green fluorescent protein (GFP) fused with a number of LCD candidates (**Figure 1B**), derived from Nup98 (Schmidt and Görlich, 2015), FIB1 (Berry et al., 2015), hnRNPDL (Batlle et al., 2020), RBM14 (Hennig et al., 2015), TAF15 (Couthouis et al., 2011), TDP-43 (Saini and Chauhan, 2011), CPEB2N (Kato et al., 2012), and hnRNPH1 (Wang et al., 2018). In addition, several optimized LCD combinations were also designed, including Hfq-GFP-FUSN, in which the weak multivalent interactions of FUSN to phase separate are enhanced by the homo-hexamer of Hfq (Murakami et al., 2015; Someya et al., 2012), and DDX4GFP, of which the DEAD-box helicase domain within DDX4 was replaced by GFP as previously described with modifications (Nott et al., 2015).

**Figure 1.**
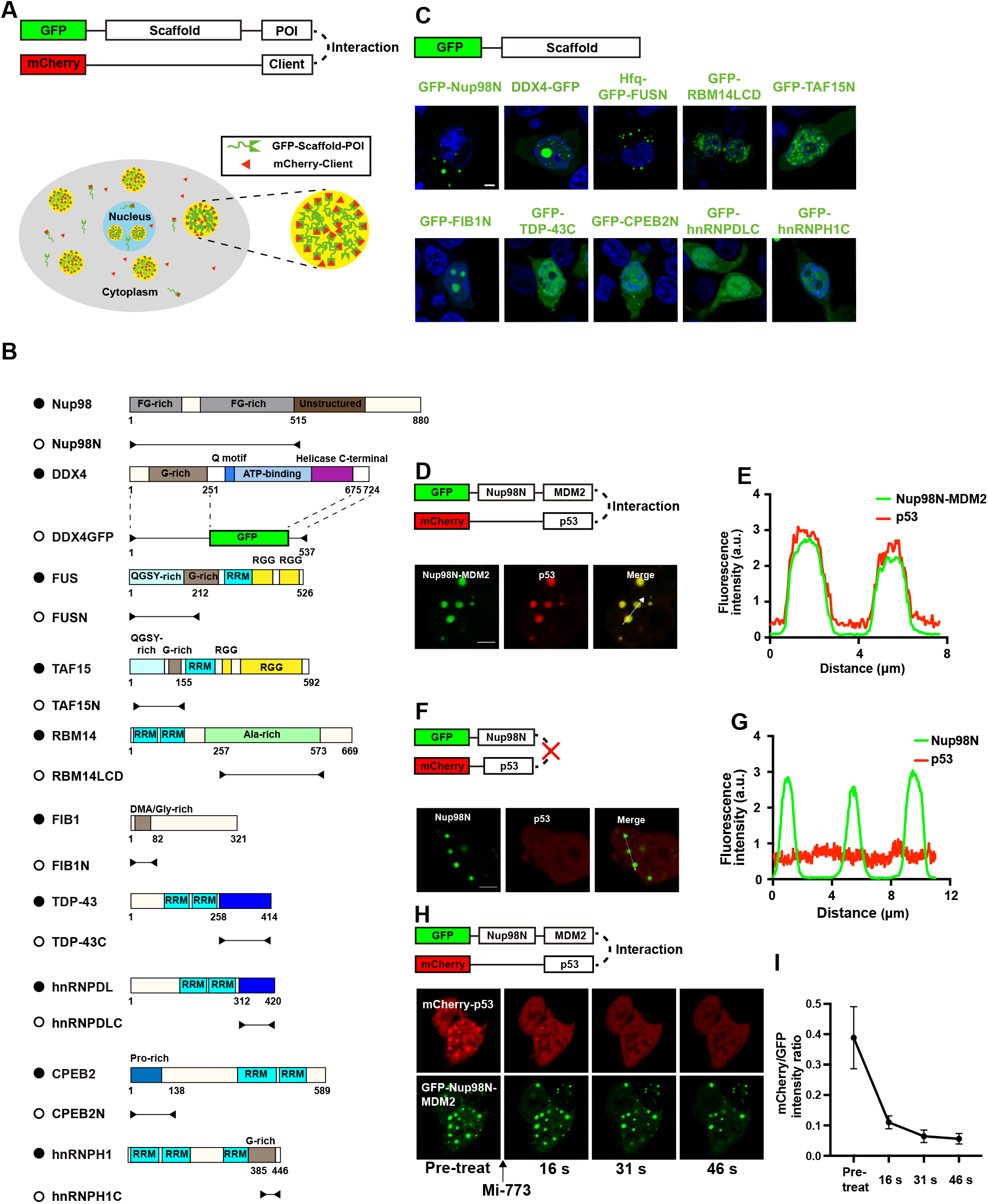
Establishment of a robust PPI-mediated recruitment system in cells. (**A**) Schematic diagram showing the strategy of CoPIC. The direct interaction between the protein of interest (POI) and the potential binding client is assessed by the enrichment of mCherry signals into the phase-separated compartments of the GFP-labeled scaffold. (**B**) The domain structures of scaffold candidates tested in CoPIC. (**C**) Characterization of scaffold candidates fused with GFP in HEK293 cells, with the nucleus stained by DAPI. Scale bars, 5 μm. (**D and F**) Validation of the direct interaction between GFP-Nup98N-MDM2 and mCherry-p53 using CoPIC. The p53 fusion protein, as indicated by the mCherry signal, was recruited to the green compartment of GFP-Nup98N-MDM2 by the specific interaction (D). The co-expression of GFP-Nup98N and mCherry-p53 served as the control (F). Scale bars, 5 μm. (**E and G**) Fluorescent intensity profiles of the lines with white arrows from D and F. (**H**) Treatment of cells co-expressing GFP-Nup98N-MDM2 and mCherry-p53 with Mi-773, an inhibitor of the MDM2/p53 interaction. Scale bars, 5 μm. (**I**) A plot of the intensity ratio of mCherry versus GFP under Mi-773 treatment.

As shown by the fluorescent images in **Figures 1C** and **S1A-S1G**, GFP-Nup98N, DDX4GFP or Hfq-GFP-FUSN formed green micron-scale membraneless spherical compartments while others exhibited poor performance of either irregularly shaped condensates or few condensates. Further analyses indicated higher sphericity of compartments of GFP-Nup98N and Hfq-GFP-FUSN (**Figure S1H**) while there was a higher proportion of spherical compartments in DDX4GFP (**Figure 1E**). All compartments formed by GFP-Nup98N, DDX4GFP or Hfq-GFP-FUSN showed fluidity when examined by fluorescence recovery after photobleaching (FRAP) assays (**Figures S1A-S1G**). Collectively, we chose GFP-Nup98N, DDX4GFP and Hfq-GFP-FUSN as potential protein scaffolds of CoPIC for further investigation.

### Nup98N condensates compartmentalize client PPI-mediated recruitment

To investigate whether the aforementioned scaffold-derived compartments support client PPI-mediated recruitment, we chose a well-known PPI pair, MDM2/p53 (Momand et al., 1992), as an example for the test. Firstly, we fused MDM2 to the C-terminus of GFP-Nup98N, DDX4GFP, or Hfq-GFP-FUSN, yielding the scaffolding vectors, GFP-Nup98N-MDM2, DDX4GFP-MDM2, or Hfq-GFP-FUSN-MDM2, respectively. Among the constructs, GFP-Nup98N-MDM2 and Hfq-GFP-FUSN-MDM2 formed robust compartments upon overexpression in HEK293 cells (**Figure 1F**) while DDX4GFP-MDM2 failed to form dynamic phase-separated compartments (**Figure S1I**). When GFP-Nup98N-MDM2 was co-expressed with the transactivation helix of p53 (15-29 aa, abbreviated to p53 hereafter) labeled with mCherry (mCherry-p53) in cells, the mCherry red fluorescence signal was enriched within the green compartments (**Figures 1D, 1E**). Moreover, the enrichment is MDM2-dependent as mCherry-p53 was not recruited into GFP-Nup98N-derived compartments (**Figures 1F, 1G**). Similar results were obtained for Hfq-GFP-FUSN-MDM2 and mCherry-p53 (**Figures S1J, S1K**).

To further confirm that the enrichment of red fluorescence signal was due to the cognate interaction between MDM2 and p53, we treated the cells co-expressing GFP-Nup98N-MDM2 and mCherry-p53 with Mi-773, a potent inhibitor specifically targeting the MDM2/p53 interaction (Wang et al., 2014). Upon treatment with Mi-773, the mCherry signal within the puncta immediately started to decrease and little remained by 46 seconds post-treatment (**Figure 1H**). Correspondingly, the ratio between the intensity of mCherry over that of GFP and Pearson’s correlation coefficient also decreased (**Figures 1I, S1L**). Notably, the rapid response to Mi-773 suggests the potential application of CoPIC in high-throughput screening of specific inhibitors for PPI, where the effectiveness of inhibitors is evaluated by the recruitment of clients into the compartments or not. Taken together, our results indicate that GFP-Nup98N can yield robust compartments for recruitment-based PPI detection in cells and thus works as the scaffold for engineered membraneless compartments used in CoPIC for subsequent studies.

### Assessment of direct and indirect interactions within the PRC2 complex using CoPIC

One of the most challenging obstacles in characterizing PPIs is to distinguish between direct and indirect interactions. Next, we evaluated whether CoPIC can fulfill the task of discriminating direct or indirect interactions. To this end, we used the well-studied polycomb repressive complex 2 (PRC2) as the testing example. PRC2 is comprised of four core subunits including enhancer of zeste 1 or 2 (EZH1/2), embryonic ectoderm development (EED), suppressor of zeste 12 (SUZ12), and retinoblastoma-binding protein 4 or 7 (RBBP4/7) (**Figure S2A**) (Margueron and Reinberg, 2011). We generated vectors expressing all PRC2 core components fused with mCherry- or GFP-Nup98N. Based on extensive biochemical and structural studies in the literature, it is known that RBBP4/7 directly interacts with SUZ12 but not with EZH2 (Glancy et al., 2020; Tan et al., 2014). We performed binary co-expression experiments to assess the interaction relationship of these components using CoPIC. Consistently, the co-expression of mCherry-RBBP4 with GFP-Nup98N-SUZ12, but not that of miRFP-RBBP7 with GFP-Nup98N-EZH2, resulted in client enrichment in the compartments (**Figures 2A, 2B, 2C and 2D**).

**Figure 2.**
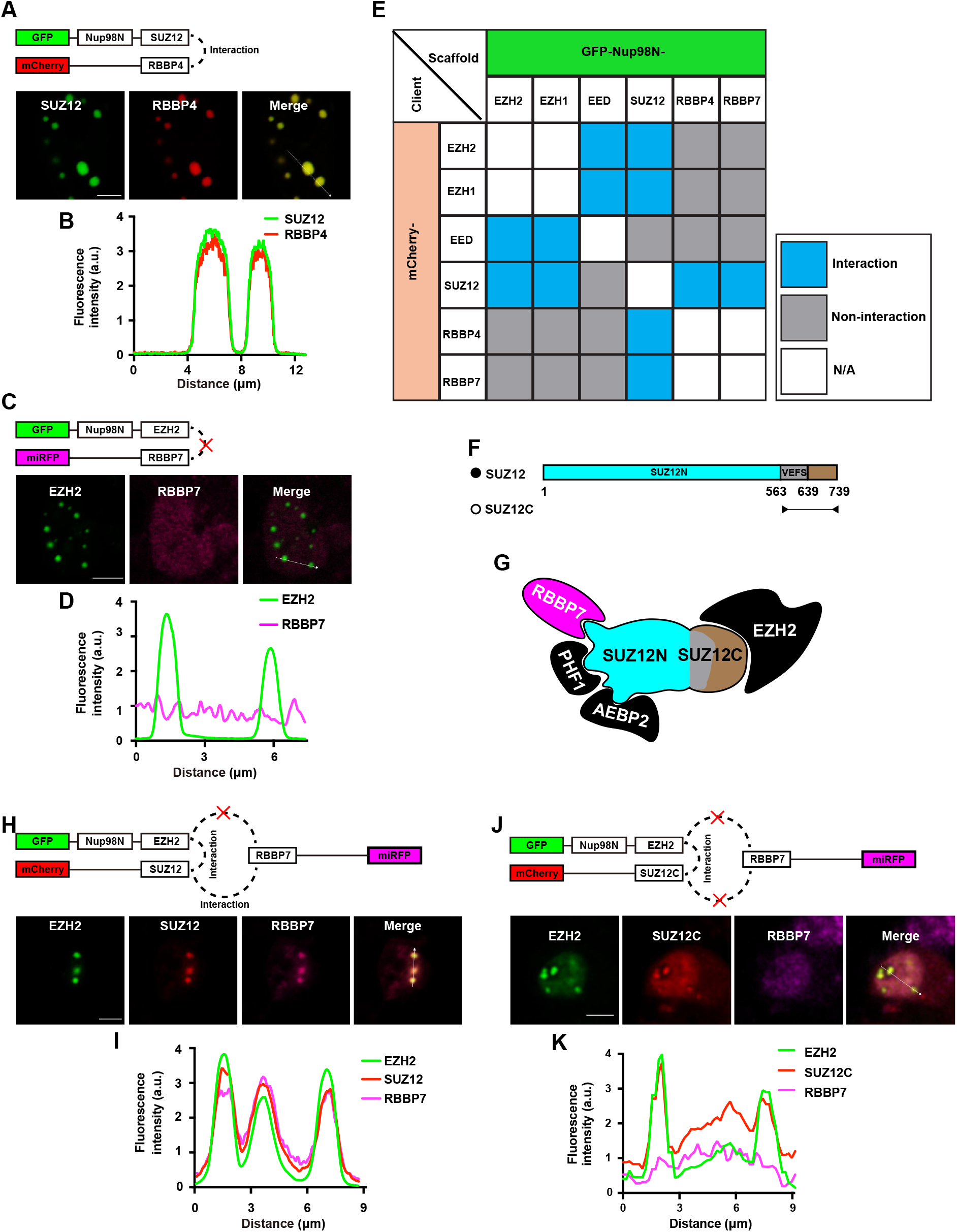
Characterization of direct and indirect interactions within the PRC2 complex using CoPIC. (**A**) CoPIC analysis of the positive interaction between SUZ12 and RBBP4. Scale bars, 5 μm. (**C**) CoPIC analysis of the negative interaction between EZH2 and RBBP7. Scale bars, 5 μm. (**E**) Summary of all pairwise interactions between PRC2 core subunits, including EZH1/2, SUZ12, EED, and RBBP4/7. (**F**) The domain structures of SUZ12 and SUZ12C (C-terminal region of SUZ12, known to bind to EZH2 but not RBBP7). (**G**) Schematic diagram showing the known factors binding to SUZ12. (**H**) CoPIC analysis of indirect interaction of EZH2 with RBBP7 mediated by SUZ12. Scale bars, 5 μm. (**J**) Verification of the bridging role of full-length SUZ12 but not SUZ12C in recruiting RBBP7 into EZH2-containing compartments. Scale bars, 5 μm. (**B, D, I and K**) Fluorescent intensity profiles of the lines with white arrows in A, C, H and J, respectively, are shown.

Further pairwise interactions between all components of PRC2 were conducted. The results were consistent with the previous studies, where the interactions between EED and EZH1/2, between EZH1/2 and SUZ12, and between SUZ12 and RBBP4/7 were direct and all other interactions were indirect (**Figure 2E**). As a key component of PRC2, SUZ12 is reported to adopt an extended conformation with the C-terminus binding EZH2 and the N-terminus binding RBBP4/7 and two accessory proteins, PHF1 and AEBP2 (**Figures 2F, 2G**). Subsequent CoPIC assays utilizing the C-terminal part of SUZ12, SUZ12C, showed that only EZH2, but not RBBP7, PHF1, or AEBP2, was enriched in the GFP-Nup98N-SUZ12C compartments (**Figure S2B**). On the contrary, CoPIC assays utilizing the full-length SUZ12 showed substantial enrichment of the signals of RBBP7, PHF1, or AEBP2 within the compartments of GFP-Nup98N-SUZ12 (**Figure S2B**).

Because we demonstrated that CoPIC can robustly detect direct PPIs, we conducted further investigation into the indirect PPIs within the PRC2 complex. Among the mutual interactions of PRC2 complex components, there are two distinctive interaction patterns: (i) direct contact of RBBP4/7 with SUZ12 mediates the indirect interaction of RBBP4/7 with EZH2; and (ii) the interaction of SUZ12 with EED requires the involvement of EZH2 (Ciferri et al., 2012; Kasinath et al., 2018). Next, we sought to characterize the typical indirect interactions via CoPIC and introduced a ternary co-expression system expressing the GFP-Nup98N-EZH2, miRFP-RBBP7, and mCherry-SUZ12. Both miRFP and mCherry signals were enriched and colocalized within the compartments of GFP-Nup98N-EZH2, indicating the bridging role of SUZ12 in recruiting RBBP7 into EZH2-containing compartments (**Figures 2H, 2I**). On the contrary, the colocalization and enrichment were not observed with co-expression of mCherry-SUZ12C (**Figures 2J, 2K**). Similar results of CoPIC also verified the indirect interaction of SUZ12 with EED via EZH2 (**Figure S2C**). Taken together, these data indicate that indirect interactions can be unambiguously demonstrated by combining binary and ternary co-transfection experiments via CoPIC.

### CoPIC screening of direct PPIs among SARS-CoV-2 RTC-related viral proteins

Taking advantage of the established CoPIC system, we screened PPIs among SARS-CoV-2 RTC-related viral proteins. The genes encoding selected viral proteins (**Table S1; Figure S3A**) were cloned into the expression vectors fused with fluorescent protein tags such as mCherry, BFP, or GFP-Nup98N. Considering the unique properties of Nsp4 and Nsp6 as transmembrane proteins and the difficulty to synthesize Nsp3, these three nonstructural proteins were not included in our analysis. First, the expression and subcellular localization of these proteins was examined in Vero E6 cells, a cell line susceptible to SARS-CoV-2 replication *in vitro* (A et al., 2020). Confocal imaging was carried out to detect the individual proteins at 24 hours post-transfection (p.t.). With adequate protein levels expressed in Vero E6 cells, representative images of cellular localization were taken with a confocal microscope. As shown in **Figure S3B**, most Nsps except Nsp1 and Nsp2 were localized both in the nucleus and cytoplasm.

After verifying the proper expression of selected viral proteins, CoPIC screening was conducted for pairwise interactions between all combinations of the 14 proteins. Among the 196 protein pairs tested, a total of 47 interaction combinations were identified as positive and summarized in a pairwise matrix (**Figure 3A**). In total, 26 interactions showed one directionality, implying a possible influence of fusion domains on the interactions. Nevertheless, 14 pairs of interactions were detected in both directions. In addition, self-interactions were observed for Nsp5, Nsp7, Nsp8, Nsp9, Nsp13, Nsp16, and N protein, suggesting that these proteins could form dimeric or multimeric complexes by interacting with themselves. However, CoPIC failed to detect the previously reported self-interactions for Nsp10 and Nsp15 (Ricagno et al., 2006; Su et al., 2006), a failure that could be due to an adverse effect of partner fusion on the self-interaction. Together, the SARS-CoV-2 RTC-related viral factors interactome-based CoPIC screening suggests concentrated interactions of open reading frame 1b (ORF1b)-encoded Nsps that were mainly connected by the Nsp12, Nsp13, Nsp14, Nsp15, and Nsp16 (**Figure 3B, 3C**). The second-most-connected proteins were Nsp5, Nsp7, Nsp8, and Nsp10 whereas none were identified for Nsp1 and Nsp2 (**Figure 3B, 3C**).

**Figure 3.**
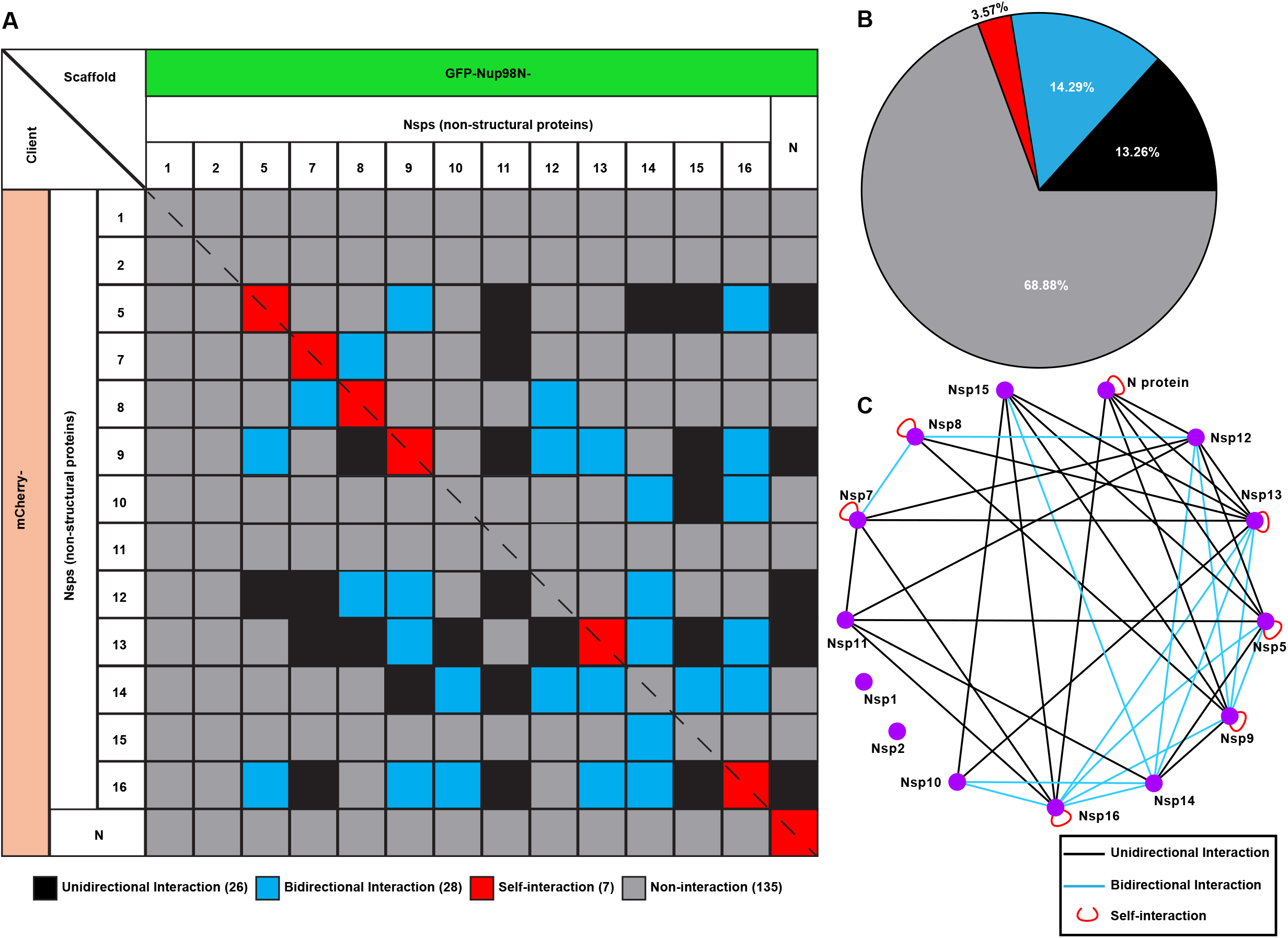
CoPIC screening of intraviral PPIs within SARS-CoV-2 RTC-related factors. (**A**) Summary of intraviral interactions between selected factors associated with RTCs. The grids with black/blue/red fill represent unidirectional/bidirectional/self-interactions identified by CoPIC while the grey ones are negative interactions. (**B**) Analysis of the proportion of each type of interaction for all pairwise interactions. (**C**) Annotation for the interaction network with degree sorted layout.

Although employing a similar expression strategy for replication, 32 novel interaction pairs in SARS-CoV-2 were identified in addition to 15 overlapped hits shared by SARS-CoV (**Figure 4A**). Among the newly discovered interaction combination in our study, co-expression of mCherry-Nsp9 and GFP-Nup98N-Nsp5 showed the enrichment of client proteins into the compartments and strong colocalization (**Figures 4F, 4G**) compared with the negative group (Nsp8-Nsp5) (**Figures 4B, 4C and 4D**). The overlapped positive pairs in SARS-CoV and SARS-CoV-2, i.e. Nsp5-Nsp12 and Nsp9-Nsp9, were also well characterized via CoPIC analyses, implying the possible critical roles for maintaining fundamental functions in the coronavirus family (**Figures 4D, 4E, 4H and 4I**). More positive pairwise interactions are exhibited in **Figures S4A-S4C**.

**Figure 4.**
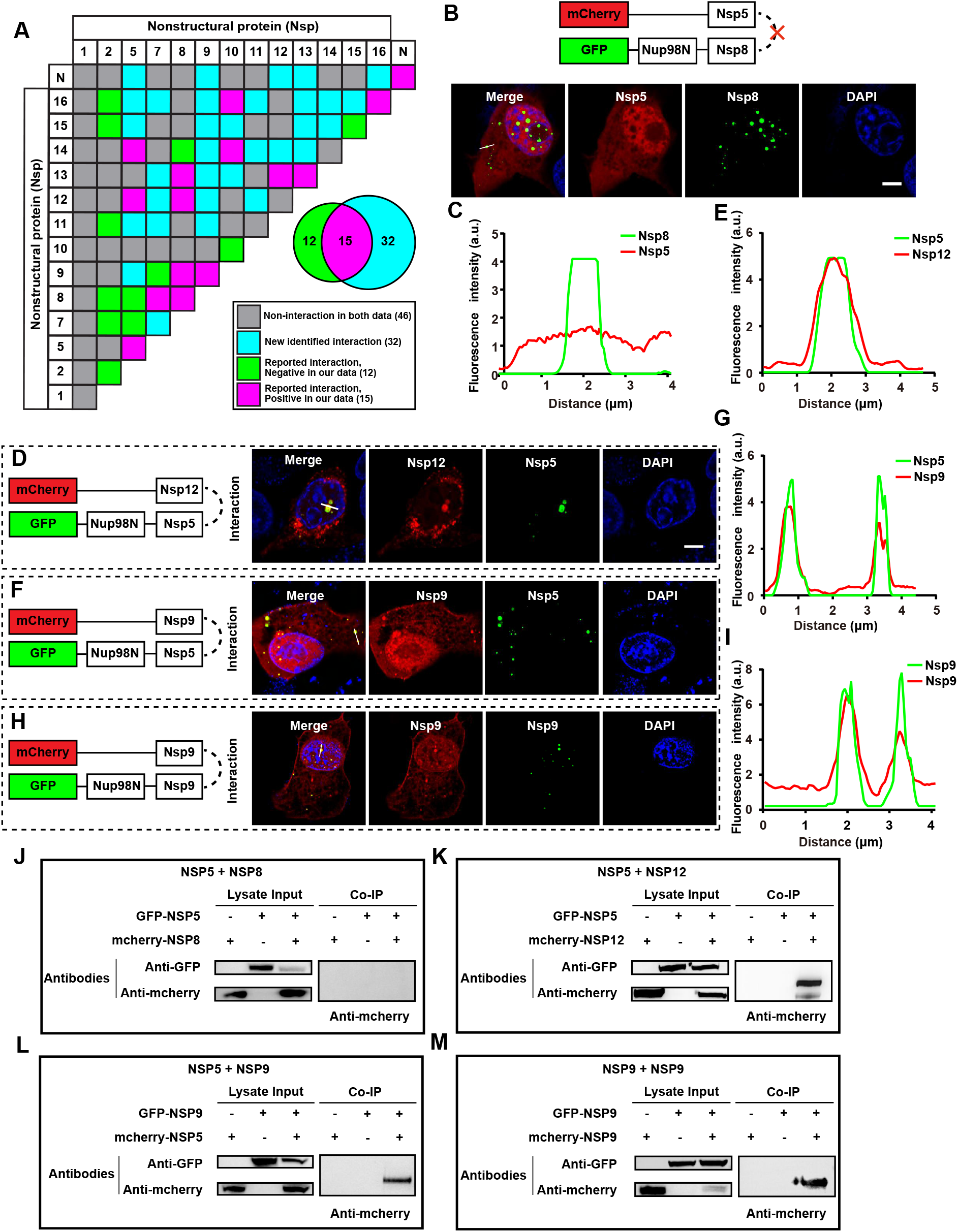
Analysis of interaction patterns between SARS-CoV and SARS-CoV-2. (**A**) Comparison of the intraviral interactions of SARS-COV2 (detected by CoPIC) with SARS-CoV (reported in the literature). The grids with magenta fill represent the interactions identified both in SARS-CoV and SARS-CoV-2; the green are the positive interactions in SARS-CoV that are negative in SARS-CoV-2 identified by CoPIC; the cyan are the novel interactions in SARS-CoV-2 identified by CoPIC; the grey are negative interactions in both SARS-CoV and SARS-CoV-2. (**B, D, F and H**) CoPIC analyses of the representative pairwise PPIs, in which Nsp5-Nsp8 are the negative interaction in SARS-CoV-2 identified by CoPIC, Nsp5-Nsp12 are well-known interactions both in SARS-CoV and SARS-CoV-2, Nsp5-Nsp9 is a novel interaction in SARS-CoV-2 identified by CoPIC, and Nsp9-Nsp9 is a self-interaction. Scale bar, 5 μm. (**C, E, G and I**) Fluorescent intensity profiles of the lines with white arrows are shown. (**J, K, L and M**) Coimmunoprecipitation analysis of the representative pairwise PPIs (B, D, F and H).

A series of coimmunoprecipitation assays were performed to validate the selected candidates. Briefly, HEK293 cells were co-transfected with two expression plasmids fused with mCherry or GFP tag. At 48 h p.t., cells were lysed and proteins were immunoprecipitated with an anti-GFP monoclonal antibody (mAb). The expression of both tagged proteins was confirmed by western blotting using respective antibodies. As demonstrated in **Figure 4J-4M**, one of the positive candidate pairs could be immunoprecipitated with another in co-expression samples but not mCherry/GFP-tagged control alone expression samples, further verifying the specific interactions among the above combinations. More pairwise interactions validated by co-IP are exhibited in **Figure S4D**.

### Characterization of in-depth interactions within SARS-CoV-2

To fulfill a variety of critical functions during the viral life cycle, viral proteins tend to form binary, ternary or larger complexes with different combinations. Considering the fact that more than half of the interaction pairs turned out to be negative, we speculated that there might be extensive indirect interactions. The corresponding interaction patterns could be discerned, and the detailed strategy of selection is as follows: for virus-encoded proteins A, B, and C, (i) A interacts with both B and C while B does not interact with C; if B could interact with C via A is involved, A, B and C are capable of forming a ternary complex (**Figure 5A, 5B**). Moreover, a large proportion of pairwise interactions with each other within multimeric complexes should also be mentioned (**Figure 5A, 5C**). Based on the above strategy, we designed the potential combinations and further investigated the detailed patterns of interactions. Among 137 potential protein combinations tested in our study, 28 positive pairs were eventually identified (**Figure 5; S5**). Taking the key pairs of Nsp7-Nsp8 as an example, there are recently published reports of structural analyses underlying the association of this combination with other Nsps including Nsp12, Nsp9, and Nsp13 (Gao et al., 2020a; Hillen et al., 2020; Wang et al., 2020; Yan et al., 2020b). However, these interactions were mainly characterized by using *in vitro* assays and have not been tested within mammalian cells. Herein, we performed CoPIC assays combined with co-IP, and further support the existence of high-order complexes. As shown in **Figure S6**, a combination of CoPIC and co-IP analyses showed a validated direct interaction between Nsp8 and Nsp12, Nsp7 and Nsp8, Nsp7 and Nsp12, thus assembling into the core complex of RTC to facilitate efficient replication. A novel ternary complex based on primase (Nsp8) and methyltransferase (Nsp16) was identified for the first time, whose formation was dependent on the involvement of Nsp7 (**Figures 6A, 6B**). The contrary results were shown in the combination of Nsp1-N-Nsp13, in which Nsp13 failed to mediate the indirect interaction between Nsp10 and N (**Figure 6C**). Collectively, extensive high-order complexes within SARS-CoV-2 were newly identified via the unique advantage of the CoPIC system, providing clues for further functional studies.

**Figure 5.**
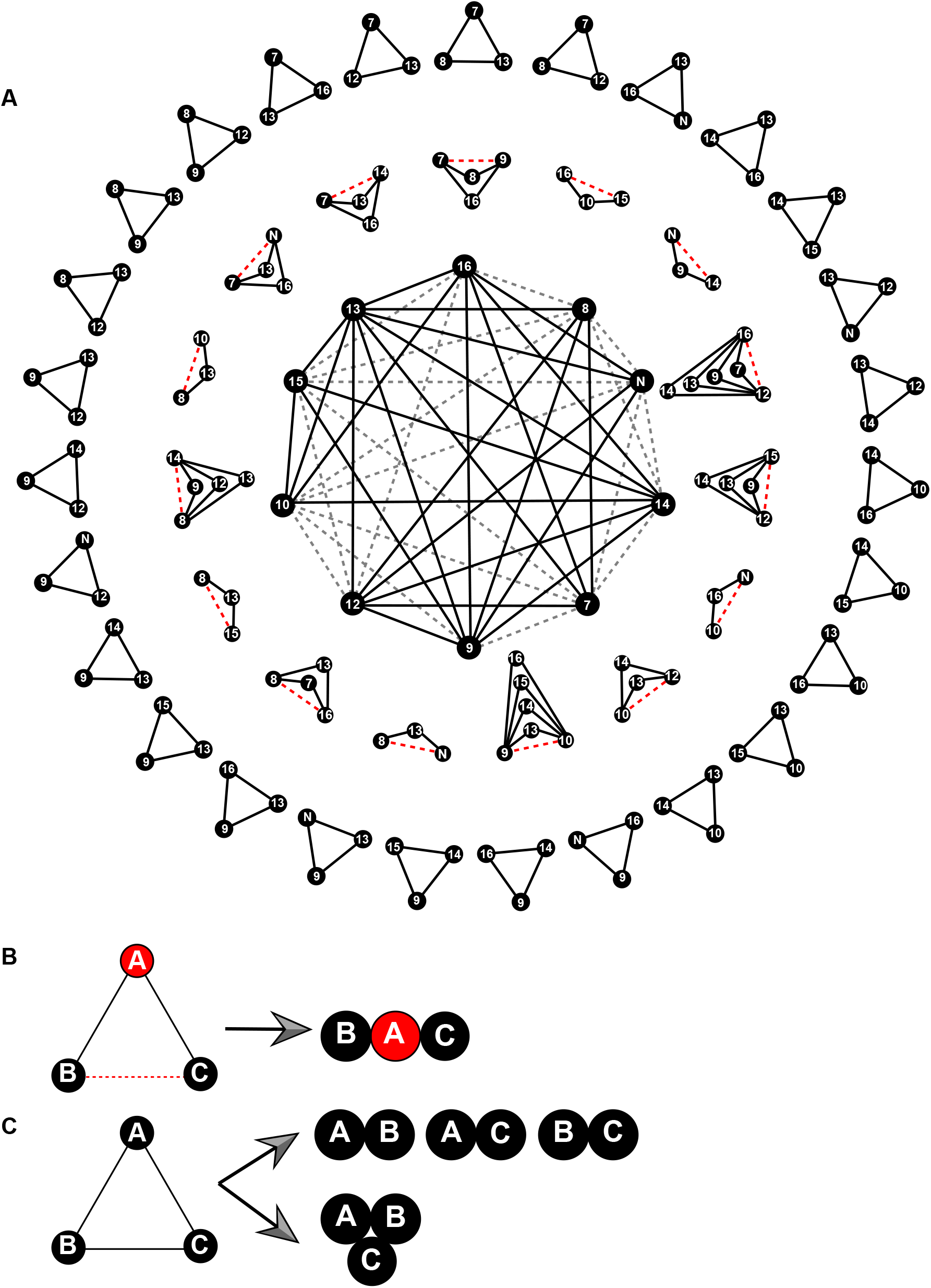
Mapping of extensive high-order interactions within SARS-CoV-2. (**A**) The network of selected viral proteins as the blueprint for mapping high-order complexes. The black solid lines are shown as the direct binary interactions while the red dashed lines are potential indirect interactions with the aid of extra intermediates. The triangles are the hypothetical or CoPIC-verified tertiary complexes and are listed in the outer ring and middle ring, respectively. (**B and C**) Interpretation of the patterns of high-order complexes. Direct contacts of A with both B and C mediate the indirect interaction of B with C, thus forming tertiary complexes shown in the middle ring (B). On the contrary, the mode in the outer ring illustrates two possibilities (C): (i) potential tertiary complex, in which all of the components can interact with each other, (ii) the combinations of multiple binary interactions.

**Figure 6.**
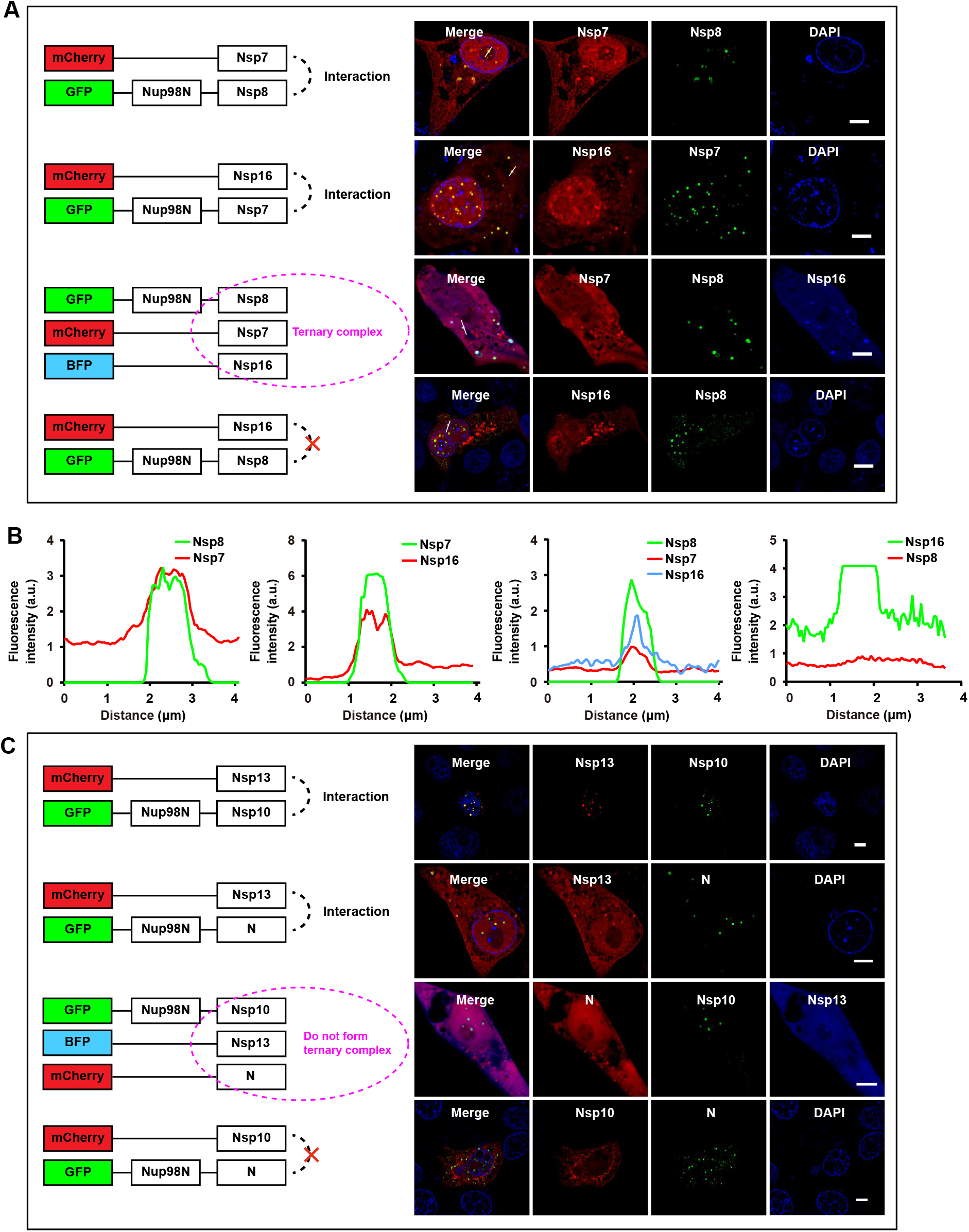
Characterization of representative indirect interactions of Nsp7-Nsp8-Nsp16 and Nsp10-N-Nsp13. (**A**) CoPIC analysis of the pairwise interactions within the positive combinations of Nsp7-Nsp8-Nsp16. Scale bar, 5 μm. (**B**) Fluorescent intensity profiles of the lines with white arrows from A. (**C**) CoPIC analysis of the pairwise interactions within the negative combinations of Nsp10-N-Nsp13. Scale bar, 5 μm.

### Analysis of the interaction hub concentrated on Nsp12-Nsp16

It is well known that ORF1ab of coronavirus is translated upon ribosomal frameshift inside ORF1a, implying significantly lower levels in producing Nsp12-Nsp16 than ORF1a-encoded products. Even so, Nsp12-Nsp16 are thought to serve as a platform to recruit their cofactors (Nsp7-Nsp10) to the central RTC, thus ensuring efficient replication and transcription. Consistent with this idea, the CoPIC screening demonstrated the concentrated interactions of Nsp12-Nsp16 with their partners (**Figures 3, 4, 5**), suggesting the potential roles of intraviral interactions in the sophisticated regulation of enzyme activities.

Taking the canonical RNA-dependent RNA polymerase as an example, Nsp12 exhibits poorly processive RNA synthesis *in vitro*, contrasting with the efficient replication of coronavirus RNA genomes with a large size *in vivo* (Ahn et al., 2012; Cheng et al., 2005; Te Velthuis et al., 2010). However, the involvement of the Nsp7/Nsp8 complex can increase the binding of Nsp12 to RNA, thus activating and conferring the processivity of Nsp12 to the RNA synthesis (Subissi et al., 2014). The Nsp7/Nsp8 complex increases the binding of Nsp12 to RNA, resulting in a larger stretch of nucleotide synthesized per binding event. Nsp8 works as RNA primase and Nsp7 might act to stabilize Nsp8 (Gao et al., 2020b; Imbert et al., 2006; Yan et al., 2020a; Zhai et al., 2005). Correspondingly, our CoPIC analysis combined with co-IP confirmed direct interactions between Nsp12 and Nsp7/8 (**Figure S6**), consistent with the newly revealed atomic structure (Hillen et al., 2020).

For another presumed RTC catalytic core, Nsp13 helicase activity is also stimulated significantly in the presence of Nsp12 by a direct interaction identified both in SARS-CoV (Adedeji et al., 2012) and our study (**Figure 3; S4**). Cryo-electron microscopy observation revealed a novel architecture of SARS-CoV-2 mini RTCs assembled by the dynamic interactions of Nsp13 with RdRp complex (Yan et al., 2020b), whose reconstruction *in vitro* was also partially verified by the CoPIC analyses in the mammalian cells (**Figure 6; S5**). Likewise, *in vitro* studies showed that the extra addition of the Nsp10 protein enhances the weak Nsp14 ExoN activity (Baddock et al., 2020; Bouvet et al., 2012). In the case of Nsp16, a strong interaction of Nsp16 with Nsp10 was shown to trigger its 2’-O-MTase activity (Bouvet et al., 2010; Imbert et al., 2008; Lin et al., 2020; Mahalapbutr et al., 2020). The direct interactions of both Nsp10-Nsp14 and Nsp10-Nsp16 were also validated by CoPIC (**Figure 3; S4**), implying the importance of Nsp10 in two distinct regulatory mechanisms (Nsp14-ExoN and Nsp16-2’-O-MTase). Moreover, the CoPIC screening identified for the first time the novel interactions between Nsp14 and Nsp16 (**Figure 3; S4**). There might be a tertiary complex of Nsp14-Nsp10-Nsp16, in which Nsp10-dependent activation of both enzymes may happen simultaneously. Nevertheless, more evidence would be required to confirm the non-limiting levels of Nsp10 to simultaneously activate the enzyme activities of Nsp14 and Nsp16.

Combined with the newly identified interaction of Nsp12 with Nsp14, an N7-MTase that can methylate the first G of viral RNA (Ferron et al., 2018; Ma et al., 2015; Yan et al., 2020a), we surmise that Nsp7/Nsp8/Nsp12/Nsp13/Nsp14 might tend to constitute a higher-order complex as the minimal viral replisome. If so, such assembly would be in charge of more capping activities, i.e., RTPase and N7-MTase, as well as unwinding of RNA secondary structures, RNA polymerization and even mismatch excision by the 3’- to-5’ exoribonuclease activity of Nsp10-Nsp14. Under the last capping step, the Nsp10/Nsp16 complex would undergo methylation of cap-0 to cap-1 through its 2’-O-MTase activity (Krafcikova et al., 2020; Rosas-Lemus et al., 2020), and might act periodically during Nsp7/Nsp8/Nsp12/Nsp13/Nsp14-mediated replication. Collectively, our interactome studies confirmed the significance and necessity of the key enzymes encoded by replicase ORF1b forming high-order complexes with the cofactors for efficient replication and transcription during the viral life cycle.

### Exploration of potential higher-order complexes

Based on the screening by CoPIC, we constructed a comprehensive high-order interactome for SARS-CoV-2 RTC-related viral proteins (**Figure 3**). The resulting map reveals a focus on ORF1b-centered interactions that are mainly connected by Nsp12, Nsp13, Nsp14, Nsp15, and Nsp16, with associated enzymatic activities that are essential for viral RNA replication and protein translation. Further extensive interactions combined with the corresponding high-order complexes imply a crucial role for Nsps in recruiting core replication/transcription proteins for the assembly of SARS-CoV-2 RTC and thus performing diverse functions during the viral life cycle. Integrated analyses of tertiary complexes identified by CoPIC revealed the various possibilities to trigger indirect interactions (**Figure 6**). Examples include the indirect interaction of Nsp7 with Nsp9 dependent on the involvement of intermediates including Nsp8, Nsp12, Nsp13, or Nsp16.

Since the coronavirus RTCs engage in a variety of RNA synthesis processes, one type of complex may be converted into another by the binding or release of specific protein partners. In this manner, such protein factors may either regulate the balance between different processes or directly form more sophisticated complexes with the core enzymes. Correspondingly, we derived several potential higher-order complexes comprising direct and indirect interactions. For example, the tandem mode of Nsp8-Nsp9-Nsp14-N illustrates the cascaded linkage of indirect interactions Nsp8-Nsp9-Nsp14 and Nsp9-Nsp14-Nsp10, in which Nsp7 and N bind to the different interfaces of the core Nsp8-Nsp9 and share no any contacts (**Figure 7A**). Multiple pairs of indirect interactions shared by the same intermediates result in a Y-shaped mode of Nsp8-Nsp10-Nsp13-N (**Figure 7B**), which emphasizes the mutual linkages of binding factors with intermediates in different interfaces with no interactive events between themselves (**Figure 7B**). On the contrary, a quinary tandem combination of Nsp7-Nsp8-Nsp9-Nsp14-Nsp10 comprising different intermediates was identified, exhibiting the successive chain (Nsp7-Nsp8-Nsp9, Nsp8-Nsp9-Nsp14, Nsp9-Nsp14-Nsp10) without forming a closed loop by end-to-end interaction (**Figure 7C**). More detailed high-order complexes are shown in **Figure S7**.

**Figure 7.**
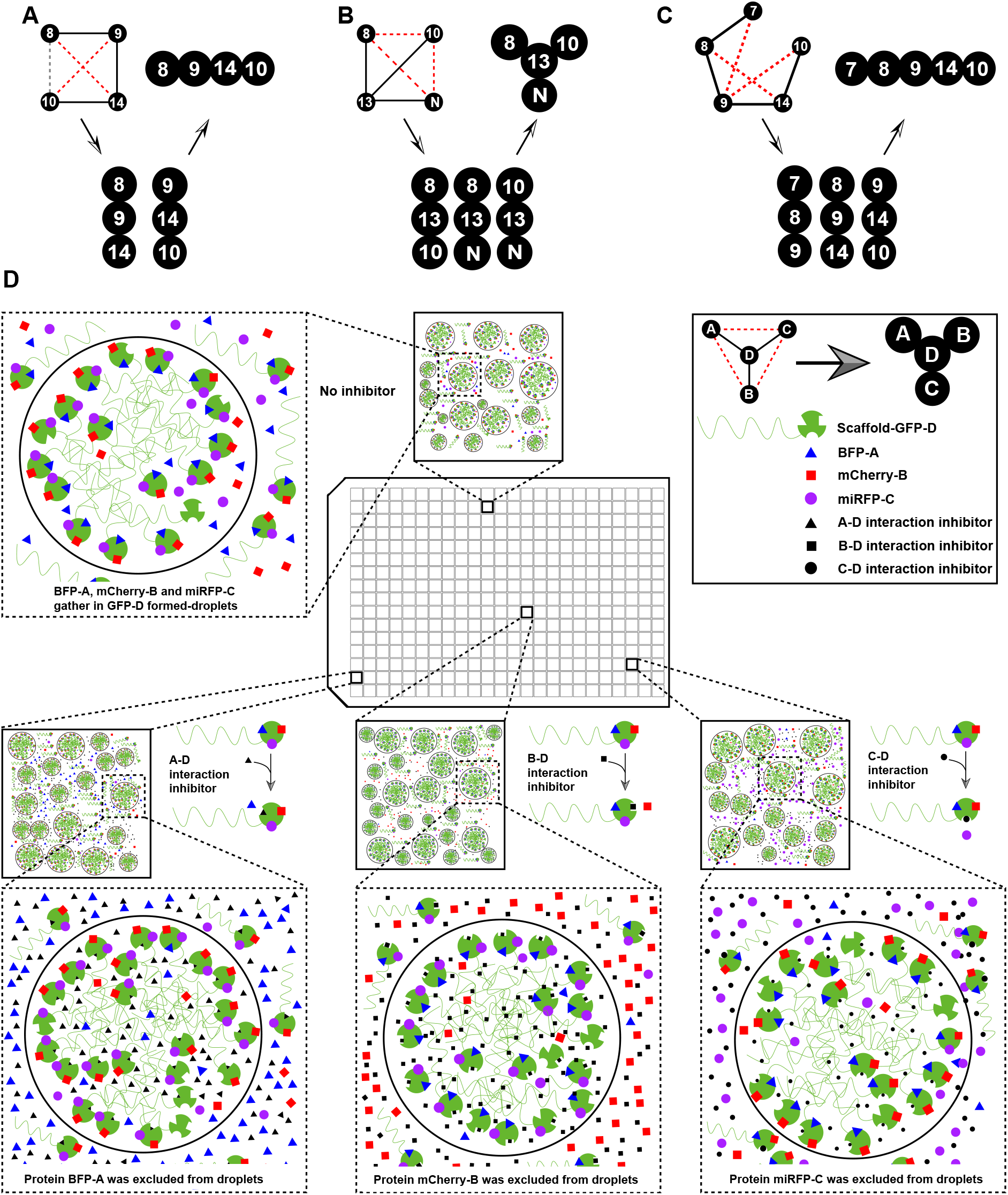
The outlook for higher-order complexes and a potential HTS scheme for PPI inhibitors. (**A, B, and C**) Simplified model systems deciphering the potential higher-order complexes. (**D**) Schematic diagram of high-throughput screening for specific inhibitors against intraviral complexes.

The above high-order complexes based on CoPIC also exhibit good potential applications in the high-throughput screening for inhibitors against PPIs (**Figure 7D**). Simply put, the core intermediate (D) fused with the scaffold acts as the platform to recruit other client components (i.e., A to C) by different interfaces and thus form the high-order complex. The subsequent screening for effective inhibitors disrupting the pairwise interactions would be performed in combination with an imaging-based method, and the activity of each candidate would be determined by assessing the enrichment of a fluorophore-labeled client into condensates enriched with its binding partner. Taken together, these simplified model systems enable more precise clues for the formation of high-order complexes during the different stages of the viral life cycle and potential therapeutic opportunities against SARS-CoV-2 by repurposing specific inhibitors to disrupt the intraviral PPIs.

## Discussion

With burgeoning knowledge about SARS-CoV-2 molecular architectures, it is essential to dissect the nonstructural protein interaction network and thus understand the assembly of RTCs. In this regard, our study established an innovative interaction screening strategy based on phase separation *in cellulo*, that is, compartmentalization of protein-protein interactions in cells (CoPIC). Since CoPIC detects the enrichment of client proteins in membraneless compartments versus the surrounding environment, discernible enrichment means the establishment of a concentration gradient. Therefore, only authentic interactions can overcome the associated entropy loss. As such, detection of nonspecific PPIs, i.e., false-positives, can be minimized. Nevertheless, if the binding partners to be recruited prefer the specific scaffold-derived condensates versus the surrounding environments, false-positive results can be generated. However, with proper controls, such false-positive hits can be detected, and a change of scaffold can be exploited to circumvent this issue.

There is another caveat associated with CoPIC. If the partner to be recruited can also undergo phase separation, the potentially complicated interaction relationship between the scaffold phase and the client phase might hinder the straightforward recruitment of the clients. For example, for the PPI screening involving the nucleocapsid protein N, there were hardly any positive interactions detected except for its self-association when N protein serves as the client; however multiple PPIs were detected when N is fused with the scaffold, GFP-Nup98N (**Figure 3A**). We reasoned that the phase separation property of N protein (Iserman et al., 2020; Perdikari et al., 2020; Zhao et al., 2020) interferes with its recruitment into the phase-separated compartment of the scaffolds. To circumvent this caveat, binding partners with phase separation propensity should be fused with the scaffold and those without phase separation propensity should serve as the clients.

Even with these caveats considered, CoPIC is a simple and efficient method for investigating PPIs in cells. In addition, CoPIC can be used to detect weak interactions since the signals are enriched within the compartments and the client proteins are overexpressed to high concentrations. Taking the interaction of Nsp14 with Nsp16 as an example, the negative pair demonstrated by the typical co-IP assay turned out to exhibit a salient bidirectional colocalization within the compartment mediated by GFP-Nup98N (**Figure 3, S4B, S4C**). Notably, CoPIC allows the compartmentalization of multiple PPIs simultaneously which is hard to achieve by other approaches and is important for the dissection of direct and indirect PPIs, hence the potential high-order complexes (Watanabe et al., 2017; Zhang et al., 2018). Better yet, an extra benefit is that CoPIC is especially suitable for PPI detection in a high-throughput fashion using an imaging-based method, e.g. high-content imaging techniques.

When applied to investigate the complicated interaction within the SARS-CoV2 proteome, the availability of CoPIC allowed the revelation of several salient findings: (i) SARS-CoV-2 shares conserved interactions with SARS-CoV and might have also evolved more novel interactions to facilitate its efficient replication/transcription; (ii) the majority of 16 Nsps are associated with viral RTC; and (iii) the interactions are centered on open reading frame 1b (ORF1b)-encoded Nsps that were mainly connected by the Nsp12, Nsp13, Nsp14 and Nsp16.

Although utilizing a similar expression strategy for efficient replication/transcription, SARS-CoV-2 exhibits a quite divergent pattern from other coronaviruses, sharing only a limited number of homologs (**Figure 4A**). By comparative analyses of the interactions between the two sub-species, SARS-CoV and SARS-CoV-2, in combination with independently published data (Li et al., 2020; Peng et al., 2008; von Brunn et al., 2007), the conserved interactions can be classified in the following categories: (i) complex formation among the scaffold proteins, where Nsp7 and Nsp8 support the RNA-dependent RNA polymerase function of nsp12, and nsp10 facilitates the methyltransferase activities of nsp14 and nsp16 by the respective associations; (ii) interaction between RNA-dependent RNA polymerase (Nsp12) and other key enzymes, including primase (Nsp8), helicase (Nsp13), and Guanine-N7 methyltransferase (Nsp14); (iii) a connection of the main protease (Nsp5) to RdRp (Nsp12) and Guanine-N7 methyltransferase (Nsp14); and (iv) self-interaction of the accessory proteins like Nsp5, Nsp7, Nsp8, Nsp9, Nsp13 and Nsp16, and structural protein N. These conserved interactions highlight the evolutionary convergence on the replication strategies at the protein-protein interaction level and point to its potentially critical role in regulating viral replication.

It is striking to identify 32 novel interactions among RTC-related factors in our study, several of which were confirmed using co-IP (**Figure 4, 5**). However, it is worth mentioning that the negative results may be due to the limitations of the assay itself and do not mean that the interaction cannot occur during infection. Of these, Nsp1 and Nsp2 have the fewest binding partners, which might result from their effect on host viability and suppressing protein synthesis (Angeletti et al., 2020; Garmashova et al., 2006; Rao et al., 2020; Yuan et al., 2020). Among the identified interactions, perhaps the most interesting are those taking place between ORF1b-encoded core enzymes (Nsp12, Nsp13, Nsp14, Nsp15, and Nsp16), responsible for the efficient genome replication and transcription. Compared with other well-known core enzymes encoded by SARS-CoV-2, the role of Nsp15 in the viral cycle is still unknown. Nsp15, also named NendoU (**<u>N</u>**idoviral **<u>endo</u>**ribonuclease, specific for **<u>U</u>**), preferentially cleaves the 3’ end of U and forms 2’-3’ cyclic phosphate ends (Kim et al., 2020). Previous reverse genetics analyses suggested the dispensable role of NendoU activity during viral replication (Ivanov et al., 2004). Mutations of the catalytic site with His and Lys residues in MHV were shown to promote a subtle defect in subgenomic mRNA accumulation while mutation of a conserved Asp residue both in HCoV-229E and MHV almost abolished viral RNA replication (Ivanov et al., 2004; Kang et al., 2007). In addition, Nsp15 might be responsible for evading the host innate immune system, which is also independent of its endonuclease activity (Liu et al., 2019; Zheng and Perlman, 2018). Intriguingly, our CoPIC screening identified multiple novel binding partners, mainly concentrated on Nsp9, Nsp10, Nsp13, Nsp14, and Nsp16, suggesting the potential involvement of Nsp15 in multiple sophisticated processes. These findings lay the foundation for further functional studies.

In summary, the current work established a sensitive method for identifying PPIs in living cells and thus constructed an interaction network of SARS-CoV-2 RTC-related factors. Among the identified interactions in combination with the data obtained by other methods, the most intriguing and striking are those between the viral core enzymes and the corresponding high-order complexes. While the functional importance of most intraviral PPIs remains less well understood, our network provides a powerful resource to dissect their roles in mediating the genome replication/transcription processes of SARS-CoV-2. The extensive interaction landscape among intraviral proteomes seems to be the norm (Bartel et al., 1996; Choi et al., 2000; Dimitrova et al., 2003; Guo et al., 2001; McCraith et al., 2000; Zell et al., 2005), underscoring the crucial roles of PPIs in the viral life cycle. Hence the disruption of these intraviral PPIs would likely provide new therapeutic opportunities against SARS-CoV-2. Notably, we have been already actively engaging in high-throughput screening for drugs disrupting essential PPIs within SARS-CoV-2 using CoPIC.

## Star⍰Methods

### Key Resources Table

#### Contact for Reagents and Resource Sharing

Further information and requests for reagents should be directed to Lead Contact Pilong Li.

#### Experimental Model and Subject Details

##### Cell lines

HEK293 cells were obtained from Cell Bank/Stem Cell Bank, Chinese Academy of Sciences (Shanghai, China), and Vero E6 cells were kindly provided by Dr. Jianyang Zeng and Dr. Dan Zhao from the Institute for Interdisciplinary Information Sciences (Tsinghua University, Beijing, China). Both HEK293 and Vero E6 cells were cultured in Dulbecco’s modified Eagle’s medium (HyClone, Inc.) supplemented with 10% fetal bovine serum (Invitrogen, Inc.) at 37°C with 5% CO_2_. FuGENE HD (Promega, Inc.) was used for transient transfection according to the manufacturer’s instructions.

#### Method Details

##### Construction of recombinant plasmids

To construct CoPIC-related vectors, the LCD regions, including the N-terminal half of Nup98 (a.a. 1-515); the N-terminal half of FIB1 (a.a. 1-82); the N-terminal half of TAF15 (a.a. 1-155); the N-terminal half of CPEB2 (a.a. 2-138); the C-terminal half of TDP-43 (a.a. 258-414); the C-terminal half of hnRNPDL (a.a. 312-420); the C-terminal half of hnRNPH1 (a.a. 385-446); the low complexity region of RBM14 (a.a. 257-573); the N-terminal half of FUS (a.a. 1-212); *Bacillus subtilis* Hfq (BsHfq); full-length (FL) EZH1; FL EZH2; FL SUZ12 (a.a. 1-739); truncated SUZ12 (a.a. 561-739); FL EED; FL RBBP4; FL RBBP7; FL PHF1, and truncated AEBP2 (a.a. 209-503), were amplified using the cDNA derived from HEK293 cells as a template and then cloned into pCDNA3.1 expression vectors. The detailed primer sequences are listed in Supplementary **Table S1**. The ORFs of SARS-CoV-2 viral proteins were synthesized *de novo* and cloned into the pCDNA3.1 plasmid by Synbio Technologies (Monmouth Junction, NJ, USA) (General Biosystems).

##### Microscopy

Imaging was with a NIKON A1 microscope equipped with a100 × oil immersion objective. NIS-Elements AR Analysis was used to analyze these images.

##### Co-immunoprecipitation assays

To confirm the interactions of viral protein pairs, Co-IP assays were performed. HEK293 cells (5×10^7^) were co-transfected with the respective expression plasmids. Cells were harvested at 48 h p.t. and washed twice with chilled phosphate-buffered saline (PBS) supplemented with protease inhibitor cocktail (Bimake, Inc.). Cell lysates were obtained using Minute™ Total Protein Extraction Kit for Animal Cultured Cells/Tissues (Invent Biotech) according to the manufacturer’s instructions. Subsequently, the lysates were incubated with GFP-Trap Magnetic Agarose (ChromoTek, Inc.) at 4°C for 1h, with rotation. After incubation, beads were collected with a Magnetic Separator, and the unbounded samples were removed and the rest saved for analysis. After 5 times washing, the samples were boiled for 10 min and subjected to SDS-polyacrylamide gel electrophoresis (PAGE) and examined by Western blot to determine if the mCherry-tagged protein was co-immunoprecipitated with the GFP-tagged protein.

##### Western blotting

All antibodies used in the study are summarized in the Key Resources Table. Figures 4J-M, S4D, and S6B were directly analyzed by western blot. The cell samples were resuspended using 2× Laemmli loading buffer. Proteins were fractionated by SDS-PAGE and transferred to a PVDF membrane. The membranes were incubated overnight with primary antibodies, then with the corresponding peroxide-labeled IgG. Finally, enhanced chemiluminescence reagent was used to visualize the results.

##### Data analysis

Images were processed with NIS-Elements AR Analysis (Nikon, Inc.) and Adobe Illustrator CC (Adobe Systems, Inc.). The fluorescence intensity was analyzed using Image J (National Institutes of Health) and the corresponding graphs were generated by Graphpad Prism8.

## Supporting information

Supplemental Key Resource Table

## Author Contributions

Conceptualization, P.L.; Methodology, H.L. and J.W; Investigation, W.X. and G.P.; Writing - Original Draft, W.X., H.L.; Writing - Review & Editing, G.P., J.W. and P.L.; Funding Acquisition, P.L., H.L.

## Acknowledgments

We acknowledge Professor Zhiyong Lou (Tsinghua University, Beijing, China) for suggestions on the manuscript. We are also grateful to SLSTU-Nikon Biological Imaging Center (Center of Pharmaceutical Technology, Tsinghua University, Beijing, China) for imaging support, and Zan Yin for illustration.

This work was supported by grants from the National Key Research and Development Program (2019YFA0508403 to P.L.), the National Natural Science Foundation of China (31800637 to J.W.; 31871443 to P.L.), and the Students Research Training Program of Tsinghua University (1911S0032 to H.L.).

## Declaration of Interests

The authors declare no conflicts of interest. The funding sponsors had no role in the design of the study; in the collection, analyses, or interpretation of data; in the writing of the manuscript; or and in the decision to publish the results.

## Figures and Table

**Table S1.**
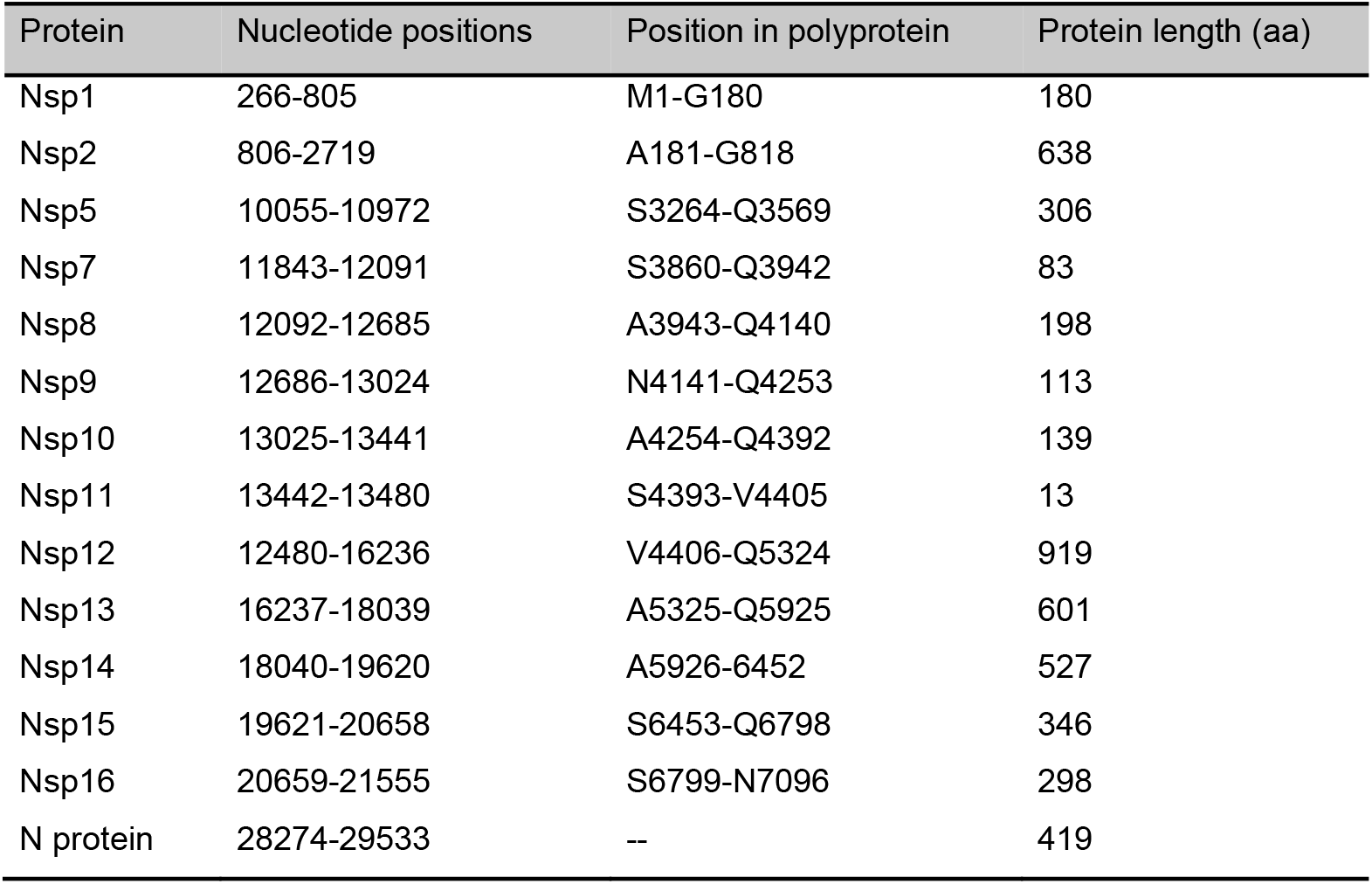
Sequence information for RTC-related viral proteins of SARS-CoV-2 used in the study for plasmid construction

## Supplementary Figures

**Figure S1. Characterization of scaffold candidates compartmentalizing and validating the direct interaction between p53 and MDM2.**

(**A-G**) FRAP analysis of scaffold candidates fused with GFP-tag in HEK293 cells. Scale bar, 5 μm.

(**H**) Histogram of the circularity of compartments formed by testing seven protein scaffolds in cells. n (Nup98N) = 100; n (DDX4GFP) = 182; n (FUSN) = 73; n (TAF15N) = 179; n (FIB1N) = 58; n (TDP-43C) = 60; n (RBM14LCD (N)) = 120. Values are means ± standard deviation.

(**I**) FRAP analysis of DDX4GFP-MDM2 in HEK293 cells. Scale bars, 5 μm.

(**J and K**) Validation of the direct interaction between Hfq-GFP-FUSN-MDM2 and mCherry-p53 using CoPIC. The p53 fusion protein, as indicated by the mCherry signal, was recruited to the green compartment of Hfq-GFP-FUSN-MDM2 by the specific interaction (J). The co-expression of Hfq-GFP-FUSN and mCherry-p53 served as the control (K). Scale bars, 5 μm.

(**L**) Plots of Pearson’s correlation coefficient for the intensity of and mCherry-p53 versus GFP-Nup98N-MDM2 pre-treatment and after 46 s.

**Figure S2. CoPIC analysis of the interaction patterns of SUZ12 with the binding factors and EZH2-mediated indirect interaction of SUZ12 with EED.**

(**A**) Schematic diagram depicting the architecture of the PRC2 complex.

(**B**) CoPIC analysis of pair-wise interactions between GFP-Nup98N fused SUZ12 (full-length SUZ12) or SUZ12C (C-terminal region of SUZ12) and mCherry-fused binding factors (EZH2, RBBP7, PHF1 and AEBP2). Scale bar, 5 μm.

(**C**) CoPIC analysis of the indirect interaction of SUZ12 with EED mediated by EZH2. Scale bars, 5 μm.

**Figure S3. Genome annotation and subcellular localization of SARS-CoV-2 RTC-related factors.**

(**A**) SARS-CoV-2 genome annotation on replication/transcription-related factors.

(**B**) Subcellular localization of SARS-CoV-2 Nsps and N protein used in CoPIC screening. Scale bar, 5 μm.

**Figure S4. Positive pairwise interactions of SARS-CoV-2 identified by CoPIC screening.**

(**A**) Collections of positive self-interactions identified by CoPIC screening.

(**B**) Collections of positive bidirectional interactions identified by CoPIC screening.

(**C**) Collections of positive unidirectional interactions identified by CoPIC screening.

(**D**) Co-IP analysis of selected pairwise interactions identified by CoPIC screening.

**Figure S5. Collections of positive tertiary complexes identified by CoPIC screening.**

**Figure S6. Characterization of mutual interactions among Nsp7, Nsp8, and Nsp12.**

(**A**) CoPIC analysis of binary and tertiary interactions among Nsp7, Nsp8, and NSp12.

(**B**) CoIP validation of pairwise interactions between Nsp7 and Nsp8, Nsp7 and Nsp12, Nsp8 and Nsp12.

**Figure S7. Higher-order complexes deduced from CoPIC analysis.**

(**A**) List of quaternary complexes deduced from CoPIC analysis.

(**B**) List of quinary complexes deduced from CoPIC analysis.

(**C**) List of six-membered complexes deduced from CoPIC analysis.

(**D**) List of seven-membered complexes deduced from CoPIC analysis.

## Notes

### Competing Interest Statement

The authors have declared no competing interest.

