## Supplemental Table 1 for "Extensive High-Order Complexes within SARS-CoV-2 Proteome Revealed by Compartmentalization-Aided Interaction Screening"

**Table S1** Sequence information for RTC-related viral proteins of SARS-CoV-2 used in the study for plasmid construction

| Protein | Nucleotide positions | Position in polyprotein | Protein length (aa) |
| --- | --- | --- | --- |
| Nsp1 | 266-805 | M1-G180 | 180 |
| Nsp2 | 806-2719 | A181-G818 | 638 |
| Nsp5 | 10055-10972 | S3264-Q3569 | 306 |
| Nsp7 | 11843-12091 | S3860-Q3942 | 83 |
| Nsp8 | 12092-12685 | A3943-Q4140 | 198 |
| Nsp9 | 12686-13024 | N4141-Q4253 | 113 |
| Nsp10 | 13025-13441 | A4254-Q4392 | 139 |
| Nsp11 | 13442-13480 | S4393-V4405 | 13 |
| Nsp12 | 12480-16236 | V4406-Q5324 | 919 |
| Nsp13 | 16237-18039 | A5325-Q5925 | 601 |
| Nsp14 | 18040-19620 | A5926-6452 | 527 |
| Nsp15 | 19621-20658 | S6453-Q6798 | 346 |
| Nsp16 | 20659-21555 | S6799-N7096 | 298 |
| N protein | 28274-29533 | -- | 419 |
