## Supplemental Key Resource Table for "Extensive High-Order Complexes within SARS-CoV-2 Proteome Revealed by Compartmentalization-Aided Interaction Screening"

**KEY RESOURCES TABLE**

| **REAGENT or RESOURCE** | **SOURCE** | **IDENTIFIER** |
| --- | --- | --- |
| **Antibodies** | | |
| mCherry (E5D8F) Rabbit mAb | Cell Signaling Technology | Cat #43590; RRID:AB_2799246 |
| GFP (D5.1) Rabbit mAb | Cell Signaling Technology | Cat #2956; RRID:AB_1196615 |
| F(ab')2-Goat anti-Rabbit IgG (H+L) Cross-Adsorbed Secondary Antibody, HRP | Thermo Fisher Scientific | Cat #A24537; RRID:AB_2536005 |
| **Bacterial and Virus Strains** | | |
| *Escherichia coli* DH5α | Tiangen Biotech | CB101-02 |
| **Chemicals, Peptides, and Recombinant Proteins** | | |
| Kanamycin | Inalco | Cat #1758-9316; CAS: 70560-51-9 |
| Ampicillin | Inalco | Cat #1758-9314; CAS: 69-52-3 |
| Phenylmethylsulfonyl fluoride (PMSF) | Amresco | Cat #0754; CAS: 329-98-6 |
| PBS | Solarbio | Cat P1020 |
| 4% Paraformaldehyde | Beyotime Biotechnology | Cat #P0099; CAS: 30525-89-4 |
| Hoechst 33258 | Beyotime Biotechnology | Cat #C1017;CAS: 23491-45-4 |
| DMEM | Hyclone | Cat #SH30022.01 |
| FBS | GIBCO | Cat #10099-141 |
| GFP-Trap Magnetic Agarose | Chromotek | Cat #gtma-100 |
| Protease inhibitor cocktail | Bimake | Cat #B14002 |
| MI-773 (SAR405838) | Selleck | Cat #S7649 |
| FuGENE^®^ HD Transfection Reagent | Promega | Cat #E2311 |
| **Critical Commercial Assays** | | |
| Total Protein Extraction Kit for Animal Cultured Cells/Tissues | Invent | Cat #SD-001 |
| BCA Protein Assay Kit | Thermo Fisher Scientific | Cat #23227 |
| Chemiluminescence reagent | Thermofisher Scientific | Cat #34077 |
| **Deposited Data** | | |
| Microscopy images | This paper |  |
| **Experimental Models: Cell Lines** | | |
| Vero E6 | J.Z. Lab | N/A |
| HEK293 | Cell Bank/Stem Cell Bank, Chinese Academy of Sciences | Cat #GNHu 43 |
| **Oligonucleotides** | | |
| GFP-NUPN (forward primer for PCR) | gtagcggcggctccctcgaaatgtttaacaaatcatttggaacacccttt | N/A |
| GFP-NUPN-3.1 (reverse primer for PCR) | cctctagatgcatgctcgagttcctcttccttcttcttagggtctga | N/A |
| GFP-NUPN-EZH1 (forward primer for PCR) | taagaagaaggaagaggaactcgagggcggtagcggtggcagcggtggtagcggcggctccatggaaataccaaatcccccta | N/A |
| GFP-NUPN-EZH1 (reverse primer for PCR) | acactatagaatagggccctctagactaaaggacgtcggtctccctctcg | N/A |
| mCherry-EZH1 (forward primer for PCR) | cggtggtagcggcggctccctcgagatggaaataccaaatccccctacct | N/A |
| mCherry-EZH1 (reverse primer for PCR) | acactatagaatagggccctctagactaaaggacgtcggtctccctctcg | N/A |
| GFP-NUPN-RBBP4 (forward primer for PCR) | taagaagaaggaagaggaactcgagggcggtagcggtggcagcggtggtagcggcggctccatggccgacaaggaagcagccttcg | N/A |
| GFP-NUPN-RBBP4 (reverse primer for PCR) | acactatagaatagggccctctagactaggacccttgtccttctggatcc | N/A |
| MCherry-RBBP4 (forward primer for PCR) | cggtggtagcggcggctccctcgagatggccgacaaggaagcagccttcg | N/A |
| MCherry-RBBP4 (reverse primer for PCR) | acactatagaatagggccctctagactaggacccttgtccttctggatcc | N/A |
| GFP-NUPN-EED (forward primer for PCR) | aagaaggaagaggaactcgagggcggtagcggtggcagcggtggtagcggcggctccatgtccgagagggaagtgtc | N/A |
| GFP-NUPN-EED (reverse primer for PCR) | cactatagaatagggccctctagatgcatgttatcgaagtcgatcccagc | N/A |
| mCherry-EED (forward primer for PCR) | aagaaggaagaggaactcgagggcggtagcggtggcagcggtggtagcggcggctccatgtccgagagggaagtgtc | N/A |
| mCherry-EED (reverse primer for PCR) | cactatagaatagggccctctagatgcatgttatcgaagtcgatcccagc | N/A |
| BFP-EED (forward primer for PCR) | aagaaggaagaggaactcgagggcggtagcggtggcagcggtggtagcggcggctccatgtccgagagggaagtgtc | N/A |
| BFP-EED (forward primer for PCR) | cactatagaatagggccctctagatgcatgttatcgaagtcgatcccagc | N/A |
| GFP-NUPN-EZH2-3.1 (forward primer for PCR) | gtagcggcggctccctcgagatgggccagactgggaagaaatctgagaag | N/A |
| GFP-NUPN-EZH2-3.1 (reverse primer for PCR) | cctctagatgcatgctcgagtcaagggatttccatttctctttcgatgcc | N/A |
| mCherry-EZH2-3.1 (forward primer for PCR) | gtagcggcggctccctcgagatgggccagactgggaagaaatctgagaag | N/A |
| mCherry-EZH2-3.1 (reverse primer for PCR) | cctctagatgcatgctcgagtcaagggatttccatttctctttcgatgcc | N/A |
| GFP-NUPN-RBBP7 (forward primer for PCR) | agaagaaggaagaggaactcgagggcggtagcggtggcagcggtggtagcggcggctccatggcgagtaaagagatgtt | N/A |
| GFP-NUPN-RBBP7 (reverse primer for PCR) | atagaatagggccctctagatgcatgttaagatccttgtccctccagttc | N/A |
| mCherry-RBBP7 (forward primer for PCR) | agaagaaggaagaggaactcgagggcggtagcggtggcagcggtggtagcggcggctccatggcgagtaaagagatgtt | N/A |
| mCherry-RBBP7 (reverse primer for PCR) | atagaatagggccctctagatgcatgttaagatccttgtccctccagttc | N/A |
| MiRFP-RBBP7 (forward primer for PCR) | agaagaaggaagaggaactcgagggcggtagcggtggcagcggtggtagcggcggctccatggcgagtaaagagatgtt | N/A |
| MiRFP-RBBP7 (reverse primer for PCR) | atagaatagggccctctagatgcatgttaagatccttgtccctccagttc | N/A |
| GFP-NUPN-SUZ12 (forward primer for PCR) | tggtagcggcggctccctcgagatggcgcctcagaagcacggcggtgggg | N/A |
| GFP-NUPN-SUZ12 (reverse primer for PCR) | ctatagaatagggccctctagatgcatgtcagagtttttgttttttgctctgt | N/A |
| mCherry-SUZ12 (forward primer for PCR) | tggtagcggcggctccctcgagatggcgcctcagaagcacggcggtgggg | N/A |
| mCherry-SUZ12 (reverse primer for PCR) | ctatagaatagggccctctagatgcatgtcagagtttttgttttttgctctgt | N/A |
| GFP-NUPN-AEBP2 (forward primer for PCR) | cggtggtagcggcggctccctcgagatgtcgtcggatggggaacccctga | N/A |
| GFP-NUPN-AEBP2 (reverse primer for PCR) | acactatagaatagggccctctagattacctcttcaacctcttctgcggc | N/A |
| mCherry-AEBP2 (forward primer for PCR) | cggtggtagcggcggctccctcgagatgtcgtcggatggggaacccctga | N/A |
| mCherry-AEBP2 (reverse primer for PCR) | acactatagaatagggccctctagattacctcttcaacctcttctgcggc | N/A |
| GFP-NUPN-PHF1 (forward primer for PCR) | cggtggtagcggcggctccctcgagatggcgcagcccccccggctgagcc | N/A |
| GFP-NUPN-PHF1 (reverse primer for PCR) | acactatagaatagggccctctagatcagaagatgccccctcctccccac | N/A |
| mCherry-PHF1 (forward primer for PCR) | cggtggtagcggcggctccctcgagatggcgcagcccccccggctgagcc | N/A |
| mCherry-PHF1 (reverse primer for PCR) | acactatagaatagggccctctagatcagaagatgccccctcctccccac | N/A |
| GFP-NUPN-SUZ12C (forward primer for PCR) | gcggtggtagcggcggctccctcgagcacaatcgtctgtatttccatagt | N/A |
| GFP-NUPN-SUZ12C (reverse primer for PCR) | gcatttaggtgacactatagaatagggccctctagatcagagtttttgttttttgctctgttt | N/A |
| mCherry-SUZ12C (forward primer for PCR) | gcggtggtagcggcggctccctcgagcacaatcgtctgtatttccatagt | N/A |
| mCherry-SUZ12C (reverse primer for PCR) | gcatttaggtgacactatagaatagggccctctagatcagagtttttgttttttgctctgttt | N/A |
| mCherry-p53 (forward primer for PCR) | gcggtggtagcggcggctccctcgagagtcaggaaacattttcag | N/A |
| mCherry-p53 (reverse primer for PCR) | cactatagaatagggccctctagagttttcaggaagtagtttcca | N/A |
| GFP-NUPN-MDM2 (forward primer for PCR) | cggtggtagcggcggctccctcgagatgactgatggtgctgtaaccacca | N/A |
| GFP-NUPN-MDM2 (reverse primer for PCR) | gacactatagaatagggccctctagagttcaccaccaccaggttgcgata | N/A |
| DDX4 (forward primer for PCR) | aatacgactcactatagggagacccaagcttgcggccgcatgggagatgaagattgggtggccgaaatcaatccgc | N/A |
| DDX4GFP 208-251-a (reverse primer for PCR) | ctccttctgcttctgacttccaagaattctttccagagcctgttattacttcttcatttaaacctttgtaaccacctcgttcacttccactgccact | N/A |
| DDX4GFP 208-251-b (forward primer for PCR) | cagaagcagaaggaggagaaagtagtgatactcaaggaccaaaagtgacctacataccccctcctccaaaagtgagcaagggcga | N/A |
| DDX4 675-724-a (reverse primer for PCR) | acccagctgtgttcaaagtgctcttgccctttctggtatcaactgatgcaaacacgtttcctcttgtactaccactgaagccaggaatgtatgtactaaaggcgtacagctcgtccat | N/A |
| DDX4 675-724-b (reverse primer for PCR) | acactatagaatagggccctctagatgcatgttactcgagatcccatgactcatcatctactggattgggagcttgtgaagaagaaaacccagctgtgttcaaa | N/A |
| GFP-CPEB2 (forward primer for PCR) | gtggtagcggcggctccctcgaaatggggccgccgccgagc | N/A |
| GFP-CPEB2 (reverse primer for PCR) | acactatagaatagggccctctagatgcatgttactcgagtcatgctgccggtacccca | N/A |
| GFP-FIBN (forward primer for PCR) | gtggtagcggcggctccctcgaaatggggatgaaaccgggc | N/A |
| GFP-FIBN (reverse primer for PCR) | acactatagaatagggccctctagatgcatgttactcgagtactctgattgccacgttt | N/A |
| GFP-hnRNPDLC (forward primer for PCR) | gtggtagcggcggctccctcgaaatggggcaacagcagcaac | N/A |
| GFP-hnRNPDLC (reverse primer for PCR) | acactatagaatagggccctctagatgcatgttactcgagtgtacggctggtaattgt | N/A |
| GFP-hnRNPH1C (forward primer for PCR) | gtggtagcggcggctccctcgaaatgggggccggtgccagc | N/A |
| GFP-hnRNPH1C (reverse primer for PCR) | acactatagaatagggccctctagatgcatgttactcgagtcatgccccagccgccgctcatac | N/A |
| GFP-RBM14LCD (forward primer for PCR) | gtggtagcggcggctccctcgaaatggggctgggcgttggc | N/A |
| GFP-RBM14LCD (reverse primer for PCR) | acactatagaatagggccctctagatgcatgttactcgagttggggtgctgttggcgtt | N/A |
| GFP-TAF15_156 (forward primer for PCR) | gtggtagcggcggctccctcgaaatggggatgagcgatag | N/A |
| GFP- TAF15_156 (reverse primer for PCR) | acactatagaatagggccctctagatgcatgttactcgagtattttcgcgctggctgtg | N/A |
| GFP-TDP-43C (forward primer for PCR) | gtggtagcggcggctccctcgaaatggggagcaatgccga | N/A |
| GFP-TDP43-C (reverse primer for PCR) | acactatagaatagggccctctagatgcatgttactcgagtattcataccccagccgct | N/A |
| Hfq (forward primer for PCR) | cccaagcttgcggccgcatgaaaccgattaacattcaggat | N/A |
| Hfq-GFP (reverse primer for PCR) | gctcactttcatggaaccgccggaaccgccttccagttccagctg | N/A |
| Hfq-GFP-FUSN (forward primer for PCR) | cggtggtagcggcggctccctcgacatggcctcaaacgattatacccaac | N/A |
| Hfq-GFP-FUSN (reverse primer for PCR) | tttaggtgacactatagaatagggccctctagatgcatgctcgaggtcctgctgtccatagccaccgctg | N/A |
| Hfq-GFP-FUSN-MDM2 (forward primer for PCR) | cggtggctatggacagcaggacctcgagatgactgatggtgctgtaacca | N/A |
| Hfq-GFP-FUSN-MDM2 (reverse primer for PCR) | tgacactatagaatagggccctctagattagttcaccaccaccaggttgc | N/A |
| mCherry-PHF1(1-511) (forward primer for PCR) | cggtggtagcggcggctccctcgagatggcgcagcccccccggctgagcc | N/A |
| mCherry-PHF1(1-511) (reverse primer for PCR) | atagaatagggccctctagatcatgagcgtctaggaagacctgg | N/A |
| **Recombinant DNA** | | |
| pCDNA3.1-mcherry-N | This paper | N/A |
| pCDNA3.1-GFP-Nup98N-N | This paper | N/A |
| pCDNA3.1-BFP-N | This paper | N/A |
| pCDNA3.1-mcherry-Nsp1 | This paper | N/A |
| pCDNA3.1-GFP-Nup98N-Nsp1 | This paper | N/A |
| pCDNA3.1-BFP-Nsp1 | This paper | N/A |
| pCDNA3.1-mcherry-Nsp2 | This paper | N/A |
| pCDNA3.1-GFP-Nup98N-Nsp2 | This paper | N/A |
| pCDNA3.1-BFP-Nsp2 | This paper | N/A |
| pCDNA3.1-mcherry-Nsp5 | This paper | N/A |
| pCDNA3.1-GFP-Nup98N-Nsp5 | This paper | N/A |
| pCDNA3.1-BFP-Nsp5 | This paper | N/A |
| pCDNA3.1-mcherry-Nsp7 | This paper | N/A |
| pCDNA3.1-GFP-Nup98N-Nsp7 | This paper | N/A |
| pCDNA3.1-BFP-Nsp7 | This paper | N/A |
| pCDNA3.1-mcherry-Nsp8 | This paper | N/A |
| pCDNA3.1-GFP-Nup98N-Nsp8 | This paper | N/A |
| pCDNA3.1-BFP-Nsp8 | This paper | N/A |
| pCDNA3.1-mcherry-Nsp9 | This paper | N/A |
| pCDNA3.1-GFP-Nup98N-Nsp9 | This paper | N/A |
| pCDNA3.1-BFP-Nsp9 | This paper | N/A |
| pCDNA3.1-mcherry-Nsp10 | This paper | N/A |
| pCDNA3.1-GFP-Nup98N-Nsp10 | This paper | N/A |
| pCDNA3.1-BFP-Nsp10 | This paper | N/A |
| pCDNA3.1-mcherry-Nsp11 | This paper | N/A |
| pCDNA3.1-GFP-Nup98N-Nsp11 | This paper | N/A |
| pCDNA3.1-BFP-Nsp11 | This paper | N/A |
| pCDNA3.1-mcherry-Nsp12 | This paper | N/A |
| pCDNA3.1-GFP-Nup98N-Nsp12 | This paper | N/A |
| pCDNA3.1-BFP-Nsp12 | This paper | N/A |
| pCDNA3.1-mcherry-Nsp13 | This paper | N/A |
| pCDNA3.1-GFP-Nup98N-Nsp13 | This paper | N/A |
| pCDNA3.1-BFP-Nsp13 | This paper | N/A |
| pCDNA3.1-mcherry-Nsp14 | This paper | N/A |
| pCDNA3.1-GFP-Nup98N-Nsp14 | This paper | N/A |
| pCDNA3.1-BFP-Nsp14 | This paper | N/A |
| pCDNA3.1-mcherry-Nsp15 | This paper | N/A |
| pCDNA3.1-GFP-Nup98N-Nsp15 | This paper | N/A |
| pCDNA3.1-BFP-Nsp15 | This paper | N/A |
| pCDNA3.1-mcherry-Nsp16 | This paper | N/A |
| pCDNA3.1-GFP-Nup98N-Nsp16 | This paper | N/A |
| pCDNA3.1-BFP-Nsp16 | This paper | N/A |
| pCDNA3.1-GFP-Nup98N | This paper | N/A |
| pCDNA3.1-DDX4-GFP | This paper | N/A |
| pCDNA3.1-Hfq-GFP-FUSN | This paper | N/A |
| pCDNA3.1-GFP-RBM14LCD | This paper | N/A |
| pCDNA3.1-GFP-TAF15N | This paper | N/A |
| pCDNA3.1-GFP-FIB1N | This paper | N/A |
| pCDNA3.1-GFP-TDP-43C | This paper | N/A |
| pCDNA3.1-GFP-CPEB2N | This paper | N/A |
| pCDNA3.1-GFP-hnRNPDLC | This paper | N/A |
| pCDNA3.1-GFP-hnRNPH1C | This paper | N/A |
| pCDNA3.1-GFP-Nup98N-EZH2 | This paper | N/A |
| pCDNA3.1-mcherry-EZH2 | This paper | N/A |
| pCDNA3.1-GFP-Nup98N-EZH1 | This paper | N/A |
| pCDNA3.1-mcherry-EZH1 | This paper | N/A |
| pCDNA3.1-GFP-Nup98N-EED | This paper | N/A |
| pCDNA3.1-mcherry-EED | This paper | N/A |
| pCDNA3.1-BFP-EED | This paper | N/A |
| pCDNA3.1-GFP-Nup98N-SUZ12 | This paper | N/A |
| pCDNA3.1-mcherry-SUZ12 | This paper | N/A |
| pCDNA3.1-mcherry-SUZ12C | This paper | N/A |
| pCDNA3.1-GFP-Nup98N-RBBP4 | This paper | N/A |
| pCDNA3.1-mcherry-RBBP4 | This paper | N/A |
| pCDNA3.1-GFP-Nup98N-RBBP7 | This paper | N/A |
| pCDNA3.1-mcherry-RBBP7 | This paper | N/A |
| pCDNA3.1-miRFP-RBBP7 | This paper | N/A |
| pCDNA3.1-GFP-Nup98N-MDM2 | This paper | N/A |
| pCDNA3.1-mcherry-p53 | This paper | N/A |
| **Software and Algorithms** | | |
| NIS-Elements AR Analysis | Nikon Corporation Healthcare Business Unit | https://www.microscope.healthcare.nikon.com/en_AOM/products/software/nis-elements |
| Adobe Illustrator CC | Adobe Systems | https://www.adobe.com/products/illustrator.html?promoid=RL89NGY7&mv=other |
| Prism 8 | GraphPad Software | https://www.graphpad.com/scientific-software/prism/ |
| Image J | National Institutes of Health | https://imagej.nih.gov/ij/download.html |
| Cytoscape | Cytoscape Software | https://cytoscape.org/ |
