## Supplemental Figure for "Extensive High-Order Complexes within SARS-CoV-2 Proteome Revealed by Compartmentalization-Aided Interaction Screening"

### Supplemeantal Figure 1

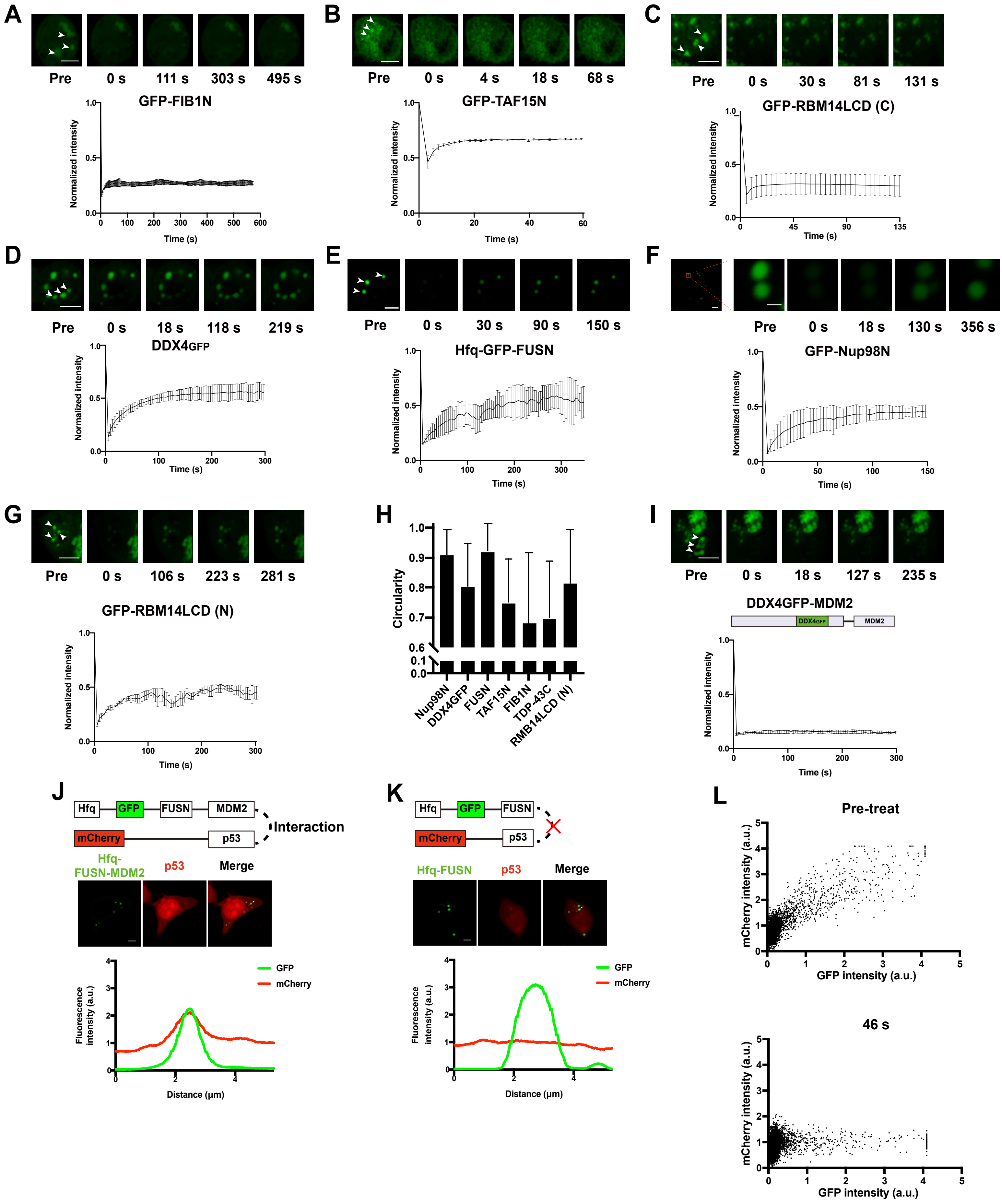

### Supplemeantal Figure 2

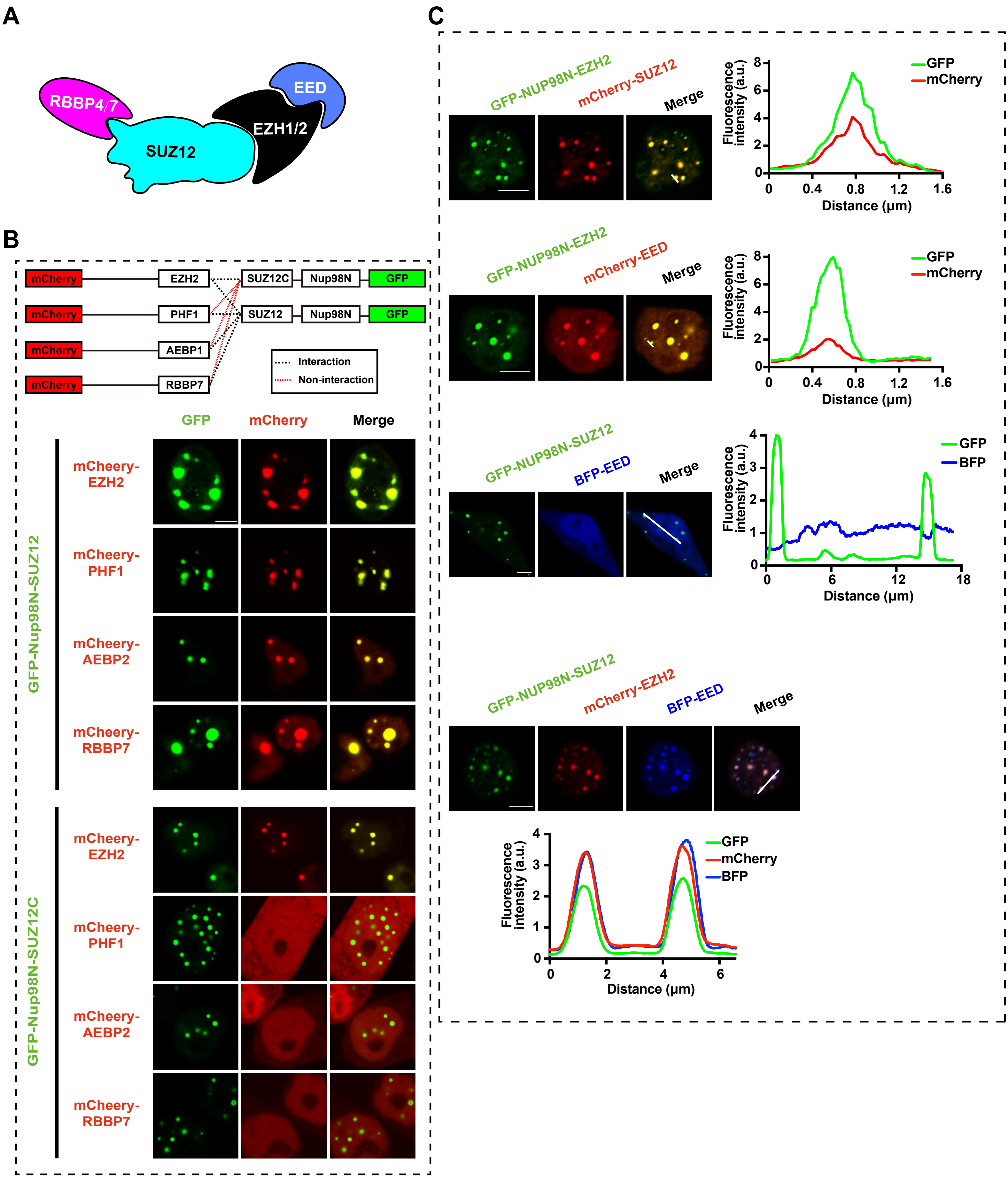

#### Supplemeantal Figure 3

A

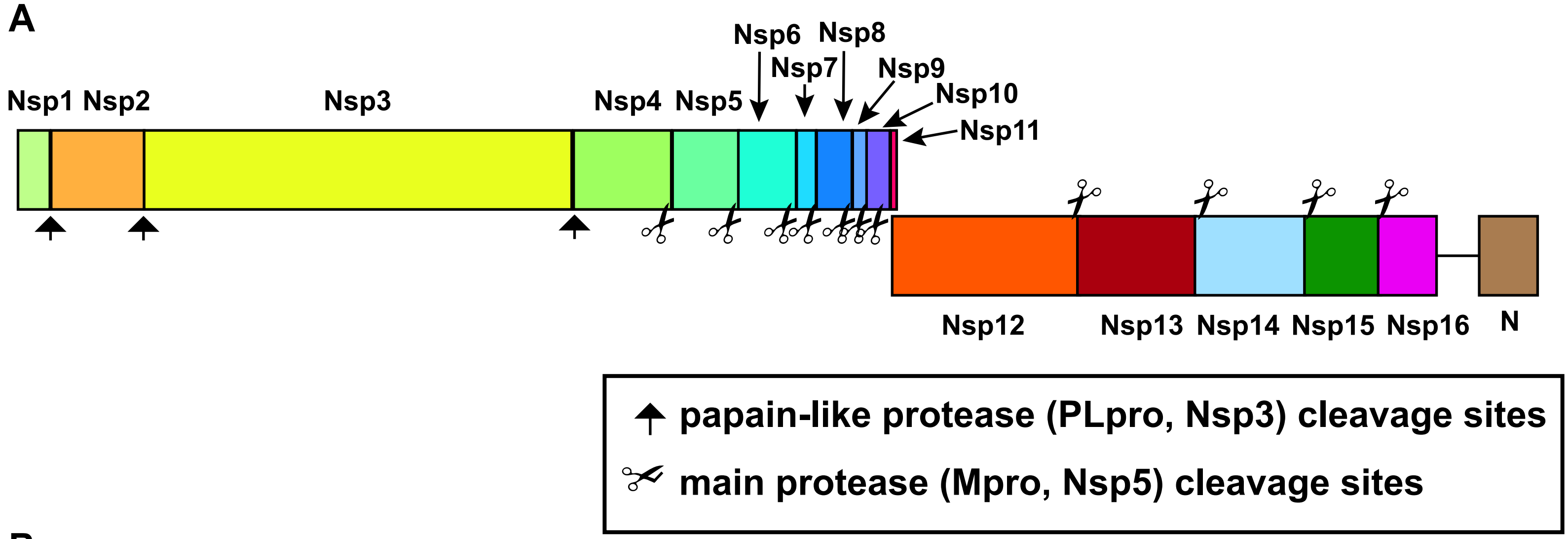


B

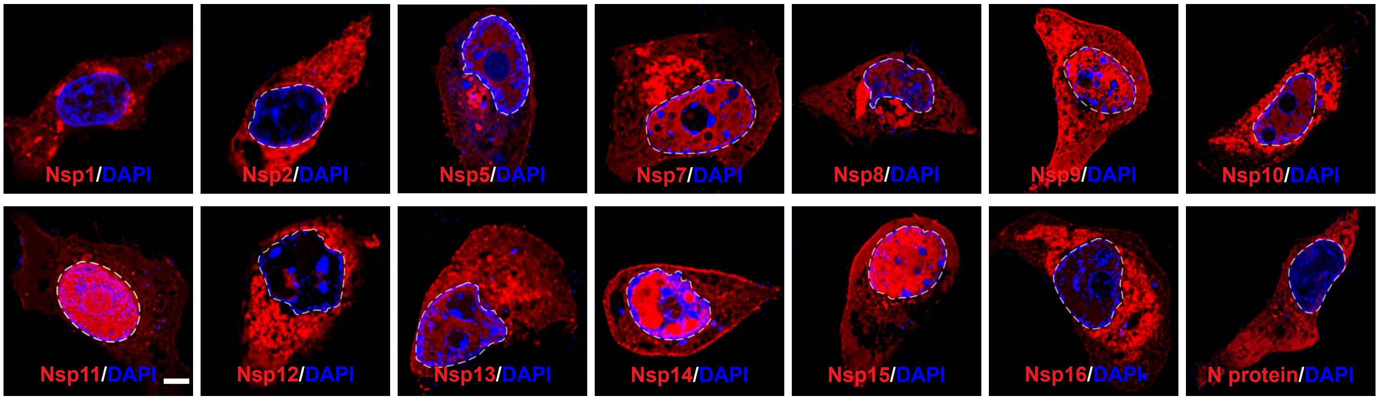

### Supplemeantal Figure 4

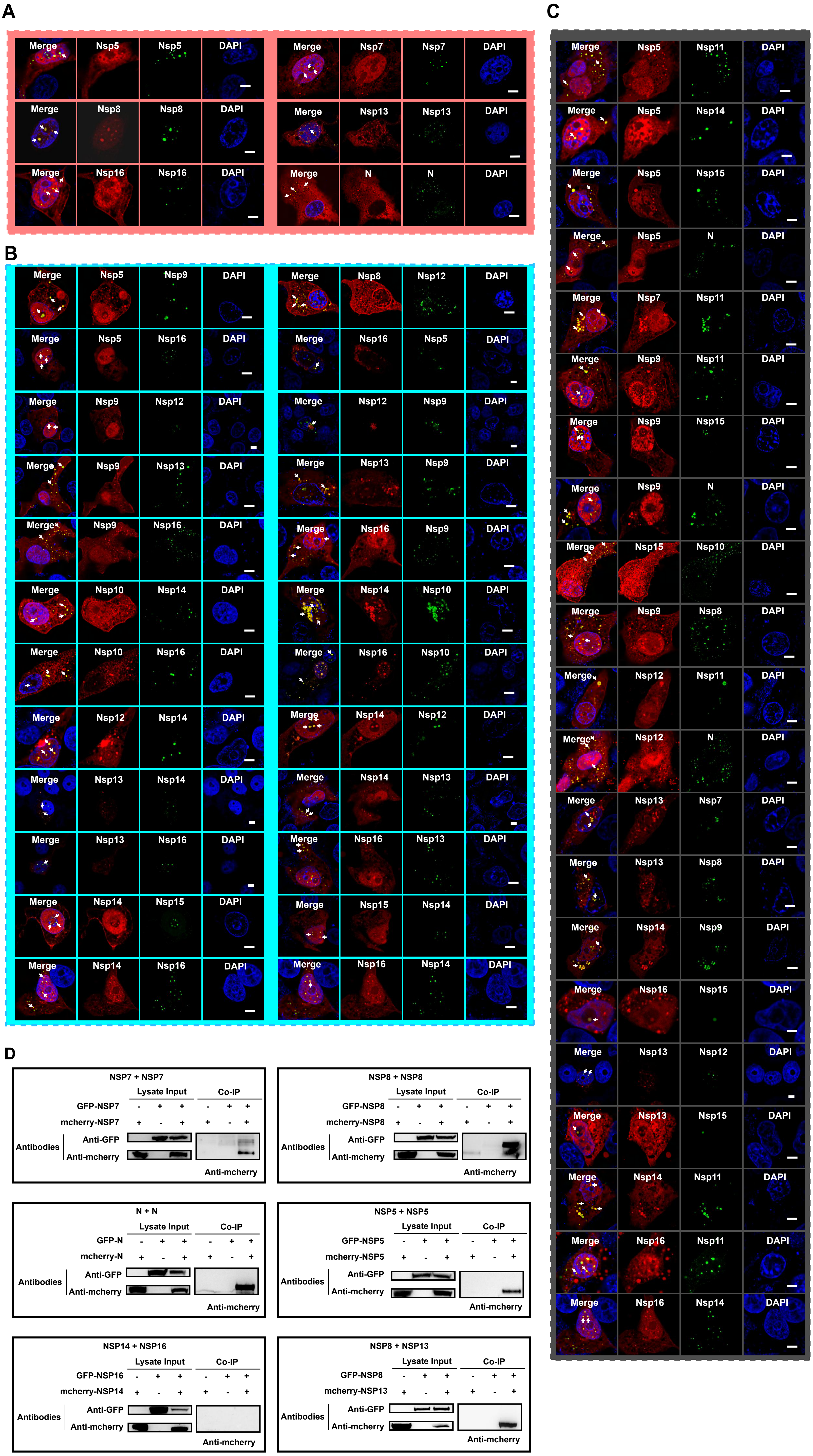

### Supplemeantal Figure 5

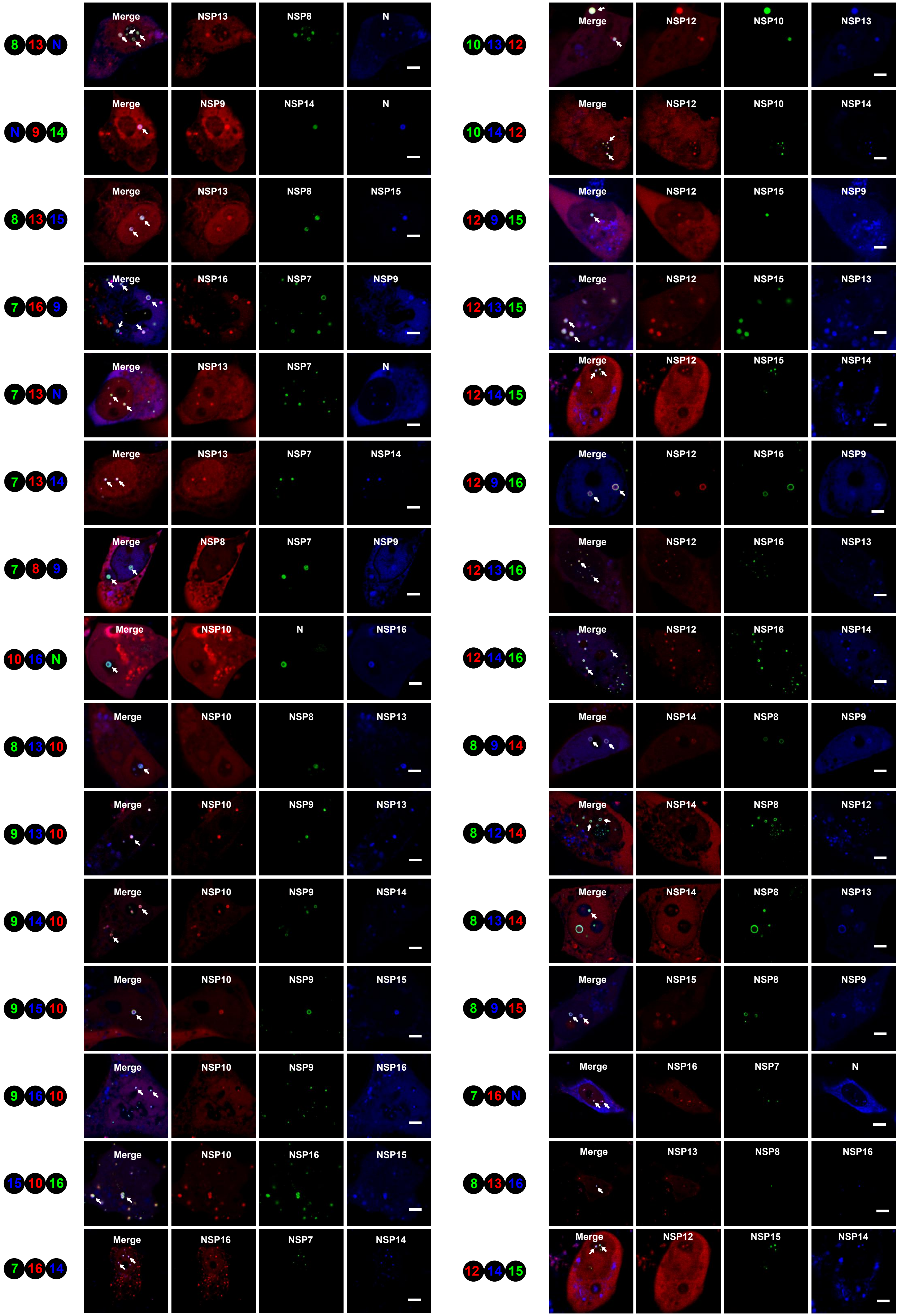

### Supplemeantal Figure 6

A

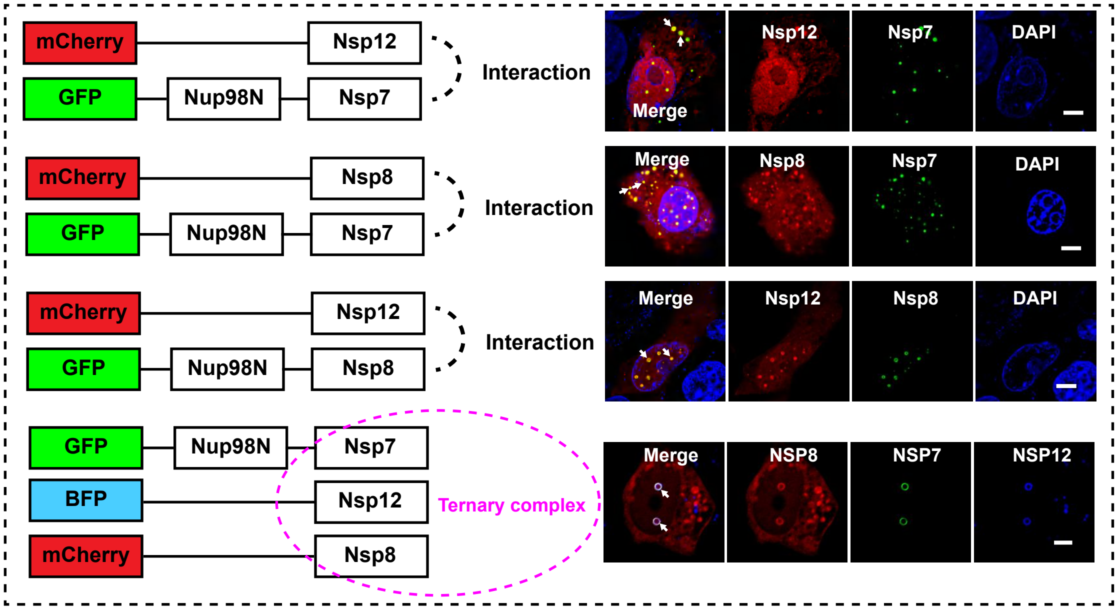


B

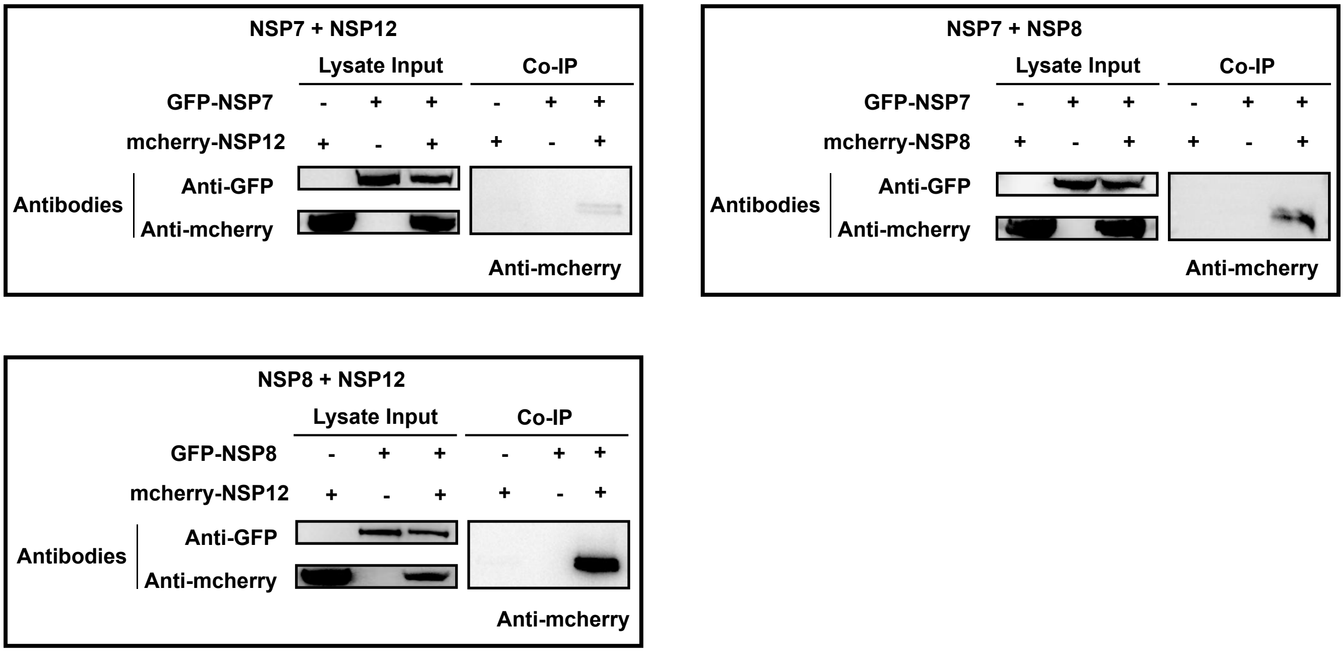

### Supplemeantal Figure 7

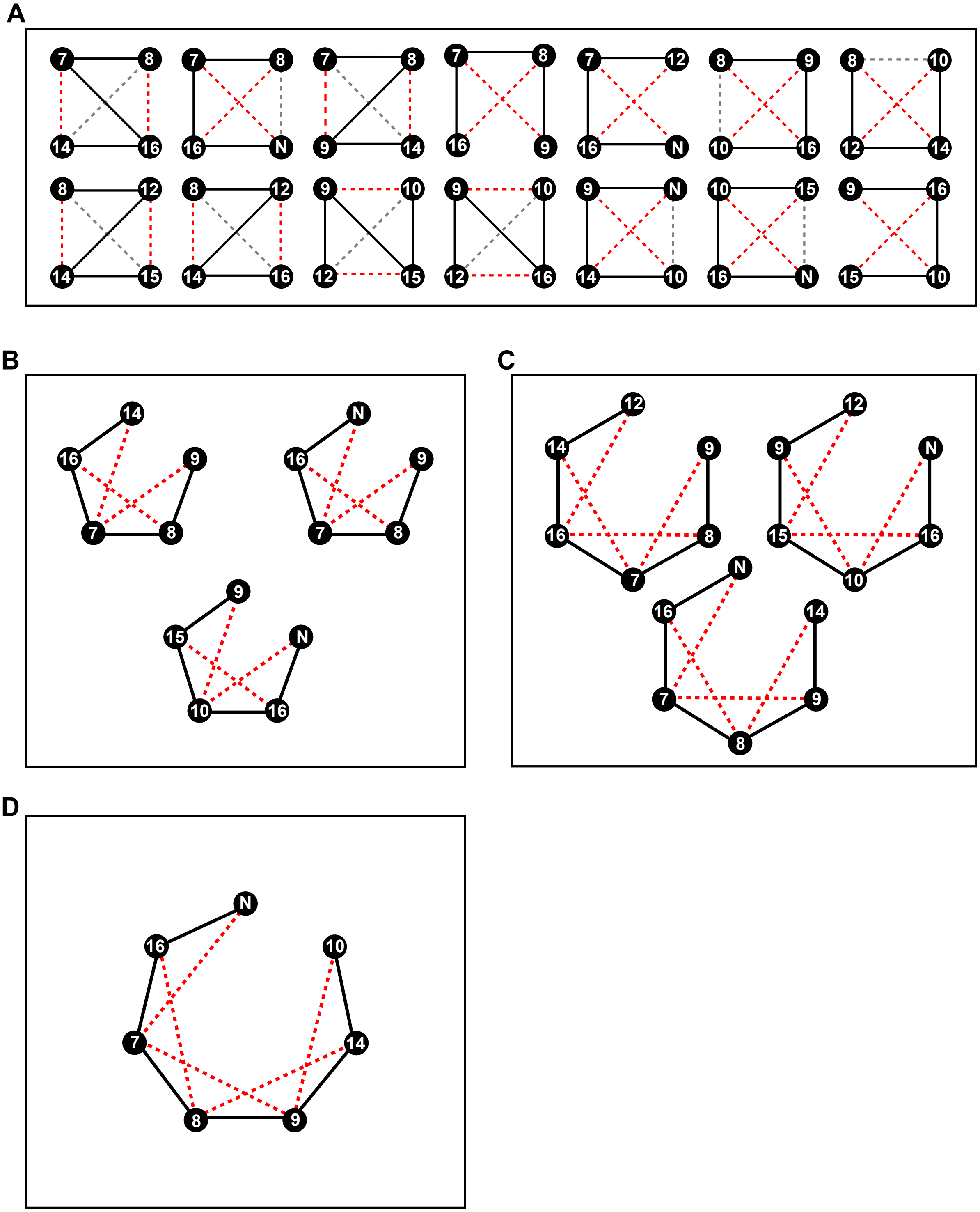
